# The allosteric modulation of Complement C5 by knob domain peptides

**DOI:** 10.1101/2020.10.24.353714

**Authors:** Alex Macpherson, Maisem Laabei, Zainab Ahdash, Melissa Graewert, James R. Birtley, Sarah Schulze, Susan Crennell, Sarah A. Robinson, Ben Holmes, Vladas Oleinikovas, Per H. Nilsson, James Snowden, Victoria Ellis, Tom Eirik Mollnes, Charlotte M. Deane, Dmitri Svergun, Alastair D.G. Lawson, Jean van den Elsen

**Affiliations:** UCB, Slough, UK. SL1 3WE; Department of Biology and Biochemistry, University of Bath, Bath, UK. BA2 7AX; European Molecular Biology Laboratory, Hamburg Unit, 22607 Hamburg, Germany; Department of Statistics, University of Oxford, Oxford, UK; Department of Chemistry and Biomedicine, Linnaeus University, 391 82 Kalmar, Sweden; Department of Immunology, Oslo University Hospital, University of Oslo, Oslo, Norway; Research Laboratory, Bodø Hospital, K.G. Jebsen TREC, University of Tromsø, Tromsø, Norway; Centre of Molecular Inflammation Research, Norwegian University of Science and Technology, Trondheim, Norway; Centre for Therapeutic Innovation, University of Bath, Bath, UK. BA2 7AX

## Abstract

To overcome limited germline combinatorial diversity, bovines have evolved a subset of antibodies with ultra-long CDRH3 regions that harbour cysteine-rich knob domains. To produce affinity-maturated peptides, we previously isolated autonomous 3-6 kDa knob domains from bovine antibodies. Here, we show that binding of four knob domain peptides elicits a range of effects on the clinically validated drug target complement C5. Allosteric mechanisms predominated, with one peptide selectively inhibiting C5 cleavage by the alternative pathway C5 convertase, revealing a targetable mechanistic difference between the classical and alternative pathway C5 convertases. Taking a hybrid biophysical approach, we present C5-knob domain co-crystal structures and, by solution methods, observed allosteric effects propagating >50 Å from the binding sites. This study expands the therapeutic scope of C5, presents new inhibitors and introduces knob domains as new, low molecular weight antibody fragments, with therapeutic potential.

---

By the end of 2019, over 60 peptide drugs have received regulatory approval, with an estimated 400 more in active development globally (Lau and Dunn, 2018; Lee *et al.*, 2019). As a potential route to discover therapeutic peptides, we previously reported a method for deriving peptides from the ultralong heavy chain complementarity determining region 3 (ul-CDRH3), which are unique to a subset of bovine antibodies(Macpherson *et al.*, 2020). We have shown that knob domains, a cysteine-rich mini-domain common to all ul-CDRH3, can bind antigen autonomously when removed from the antibody scaffold(Macpherson *et al.*, 2020). This allows peptide affinity maturation to be performed *in vivo,* harnessing the cow’s immune system to produce peptides with complex stabilising networks of disulphide bonds.

For the discovery of knob domain peptides, immunisation of cattle is followed by cell sorting of B-cells using fluorescently labelled antigen. A library of antigen-specific CDRH3 sequences is created by performing a reverse transcription polymerase chain reaction (RT PCR) on the B-cell lysate, followed by a polymerase chain reaction (PCR) using primers specific to the conserved framework regions which flank CDRH3(Macpherson *et al.*, 2020). Upon sequencing, ul-CDRH3 are immediately evident and the knob domains can be expressed recombinantly as cleavable fusion proteins(Macpherson *et al.*, 2020).

This method for discovery of knob domain peptides was established using Complement component C5, and we reported peptides which bound C5 with affinities in the pM - low nM range(Macpherson *et al.*, 2020). Herein, we use these novel peptides to probe the structural and functional aspects of C5 activation.

C5 is the éminence grise of the complement cascade’s druggable proteins, and the target of effective therapies for diseases with pathogenic complement dysregulation, of which paroxysmal nocturnal haemoglobinuria(Rother *et al.*, 2007) and atypical haemolytic uremic syndrome(Nurnberger *et al.*, 2009) are notable examples. Six monoclonal antibodies targeting C5 have reached, or are entering, clinical trials, closely followed by C5-targeting immune evasion molecules(Romay-Penabad *et al.*, 2014), aptamers(Biesecker *et al.*, 1999), cyclic peptides(Ricardo *et al.*, 2014), interfering RNA(Borodovsky *et al.*, 2014), and small molecules(Jendza *et al.*, 2019). Currently, C5 inhibitors are being trialled for the treatment of acute respiratory distress syndrome arising from SARS-CoV-2 infection(Smith *et al.*, 2020; Wilkinson *et al.*, 2020; Zelek *et al.*, 2020).

C5 is the principal effector of the terminal portion of the complement cascade. At high local C3b concentrations, arising from activation of either or both of the classical (CP) and mannose binding lectin (LP) pathways, aided by the amplificatory alternative pathway (AP), C5 is cleaved into two moieties with distinct biological functions. Cleavage is performed by two convertases; C4bC2aC3b, formed in response to CP or MBL activation (henceforth the CP C5 convertase), and C3bBbC3b, formed in response to AP activation (henceforth the AP C5 convertase). Although the constitutive components of the C5 convertases differ, they are thought to be mechanistically identical. Once cleaved, the C5a fragment is the most proinflammatory anaphylatoxin derived from the complement cascade. When signalling through C5aR1 and C5aR2, C5a is a strong chemoattractant recruiting neutrophils, eosinophils, monocytes, and T lymphocytes to sites of complement activation, whereupon it activates phagocytic cells, prompting degranulation. C5b meanwhile, interacts with C6, recruiting C7-C9 to form the terminal C5b-9 complement complex (TCC). Once inserted into a cell membrane, the TCC is referred to as the membrane attack complex, a membrane-spanning pore which can lyse sensitive cells.

Aspects of the structural biology of C5 are well understood, due to a crystal structure of the apo form(Fredslund *et al.*, 2008) and a number of co-crystal structures of C5 with various modulators. By virtue of its constitutive role in the terminal pathway, C5 is a recurrent target for immune evasion molecules and structures have been solved of C5 in complex with an inhibitory molecule derived from *Staphylococcus aureus,* SSL-7(Laursen *et al.*, 2010), as well as several structurally distinct examples from ticks: OmCI(Jore *et al.*, 2016), RaCI(Jore *et al.*, 2016) and Cirp-T(Reichhardt *et al.*, 2020). Additionally, the structures of C5 with the inhibitory monoclonal antibody (mAb) eculizumab(Schatz-Jakobsen *et al.*, 2016), of C5 with a small molecule inhibitor(Jendza *et al.*, 2019), and of C5 with the complement depleting agent Cobra Venom Factor (CVF)(Laursen *et al.*, 2011), have all been determined.

Here, we probe C5 with knob domain peptides and explore the molecular processes which underpin allosteric modulation of this important drug target. This study is the first to investigate the molecular mechanisms and pharmacology of this recently isolated class of peptide.

## Results

### Functional characterisation of anti-C5 bovine knob domain peptides

Complement activation assays were performed, using two ELISA kits which measured CP and AP activation in human serum, through C5b neo-epitope formation and C5a release. Orthogonal ELISA assays, which measured C3b and C9 deposition, were also developed (Figure 1, a-b). We tested four knob domain peptides: K8, K57, K92 and K149, which have been previously reported to display tight binding to C5(Macpherson *et al.*, 2020).

**Figure 1.**
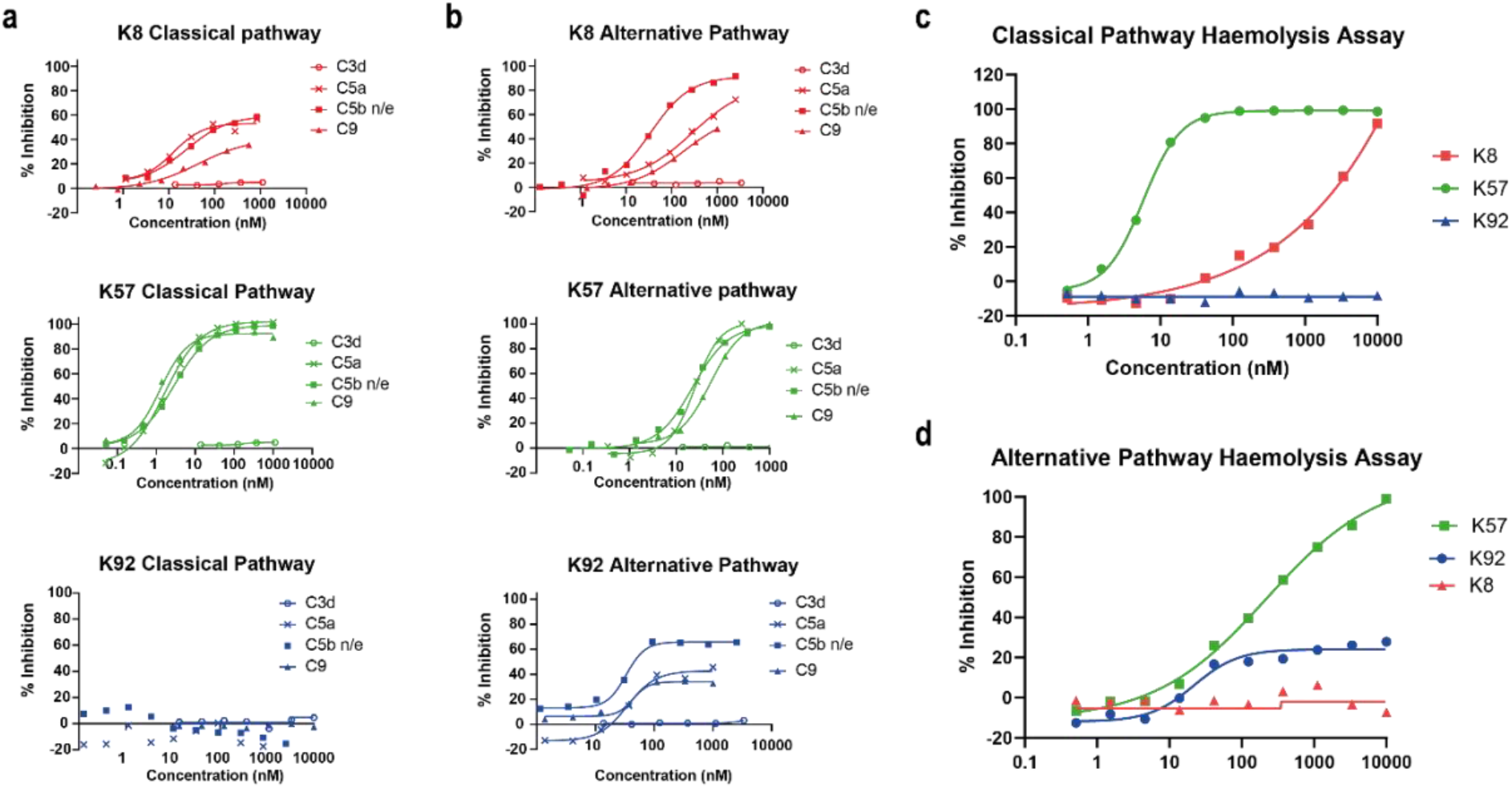
Functional modulation of C5 via knob domain peptides. Classical pathway (CP) driven ELISA assays (a) and alternative pathway (AP) driven ELISA assays (b) are shown. For both pathways, the inhibition of 3d deposition, C5b neo-epitope formation, C5a release and C9 deposition were tracked. Haemolysis assays with sheep erythrocytes, for the CP (c), and rabbit erythrocytes, for the AP (d), show that K57 is a potent and efficacious inhibitor of both pathways. K92 is selective, partial antagonist of the AP, while K8 is a weak antagonist of the CP but did not show efficacy in the AP haemolysis assay, below 10 μM.

By ELISA, K57 was a potent and fully efficacious inhibitor of C5 activation, preventing release of C5a, and deposition of C5b and C9. As expected, there was no effect on C3b, which is upstream of C5. In contrast, K149 was a high affinity silent binder with no discernible effect on C5a release, formation of C5b neo-epitope or C9 deposition, even at peptide concentrations in excess of 100 x K_D_ (Supplementary Section 1).

K8 and K92 exerted more nuanced allosteric effects on C5. By ELISA, K92 partially prevented C5 activation by the AP but, intriguingly, no effect was observed in assays where the AP component was not isolated, suggesting K92 selectively inhibits C5 activation by the AP C5 convertase. Partial antagonists, where the degree of inhibition for the asymptotic concentrations of a dose response curve (E_max_) is below 100 %, are an impossible mode of pharmacology for orthosteric antagonists(Klein, Vinson and Niswender, 2013) and we therefore propose that K92 operates by a non-steric mechanism. K8 was also demonstrably allosteric, partially inhibiting both the AP and CP in ELISA experiments. For K8 and K92, no effect on C3b deposition was detected.

When tested in CP and AP haemolysis assays, K57 was a potent and fully efficacious inhibitor of complement mediated cell lysis. Consistent with the ELISA data, K92 was active solely in the AP-driven haemolysis assay, achieving E_max_ values of 30-40 %; while K8 was efficacious in the CP assay but did not show activity in the AP assay below 10 μM, potentially a consequence of the increased serum concentration and stringency of the haemolysis endpoint.

### Crystal structure of the C5-K8 peptide complex

To elucidate the structural basis for the allosteric modulation of C5, we determined the crystal structure of the C5-K8 complex at a resolution of 2.3 Å (Supplementary 2.1 shows data collection and structure refinement statistics). The structure of the C5-K8 complex shows the K8 peptide binding to a previously unrecognised regulatory site on C5; the macroglobulin (MG) 8 domain of the α-chain (Figure 2a-b). K8 adopts a cysteine knot configuration, where a flattened 3-strand β-sheet topology is constrained by three disulphide bonds (Figure 3 and Supplementary 2.5). Analysis of the K8-C5 complex with the macromolecular interfaces analysis tool PDBePISA(Krissinel and Henrick, 2007), reveals a large interaction surface (comprising 1639Å^2^; 849Å^2^ contributed by K8 and 790Å^2^ by C5), comparable to those seen in Fab-antigen complexes, stabilised by an extensive network of 18 hydrogen bonds between K8 and the MG8 domain (Supplementary 2.6), dominated by arginine residues R23_K8_, R32_K8_ and R45_K8_. The extensive H-bond network is further bolstered by ionic interactions, between R32_K8_ and D1471_C5_ (C5 numbering based on mature sequence), D25_K8_ and K1409_C5_, and H36_K8_ and D1382_C5_ (Supplementary 2.7). The opposing face of K8 was fortuitously stabilised by a substantial, 1275.1 Å^2^, crystal contact with the C5d domain of a symmetry-related C5 molecule, (Supplementary 2.2), ensuring clear electron density and clearly enunciating the disulphide bond arrangement and backbone and side chains interactions. A mFo-DFc simulated annealing omit map of the C5-K8 complex is displayed in Supplementary 2.4, showing clear electron density for the K8 peptide. Despite the overall resolution of the dataset comparing favourably with other C5 structures in the PDB(Schatz-Jakobsen *et al.*, 2016; Laursen *et al.*, 2010; Jore *et al.*, 2016; Fredslund *et al.*, 2008), density for the C345C domain was largely absent, due to this flexible domain occupying a solvent channel.

**Figure 2.**
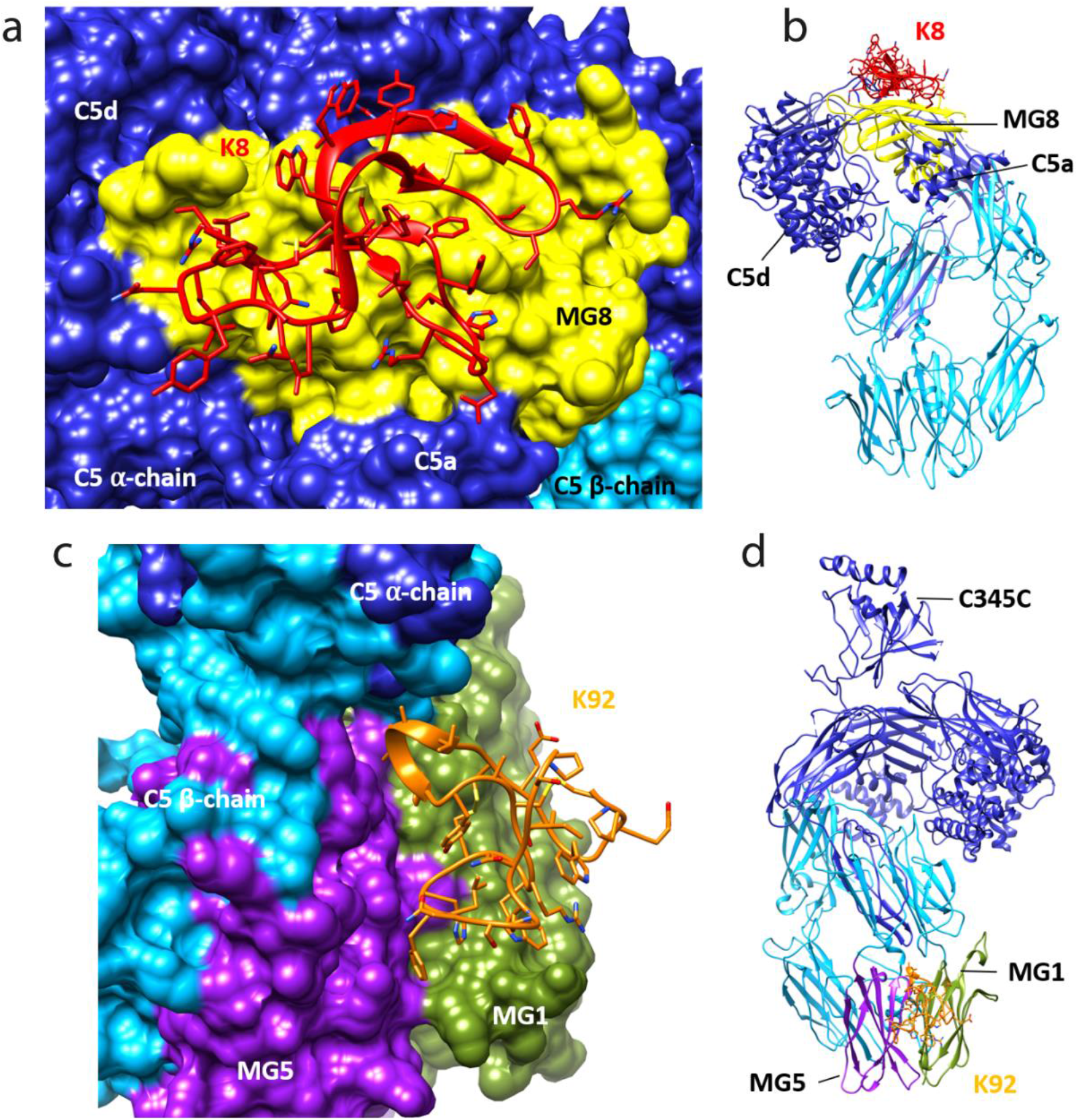
Knob domain crystal structures. Panels a and b show the crystal structure of C5 in complex with the K8 knob domain peptide. The K8 epitope is located on a previously unreported ligand binding site on the MG8 domain of C5 (a). The entire structure is shown (b), notably, electron density for the flexible C345C domain of C5 was absent, preventing building of this domain in the structure. The crystal structure of C5 in complex with the K92 knob domain peptide is shown (panels c and d). The K92 epitope is located between the MG1 and MG5 domains.

**Figure 3.**
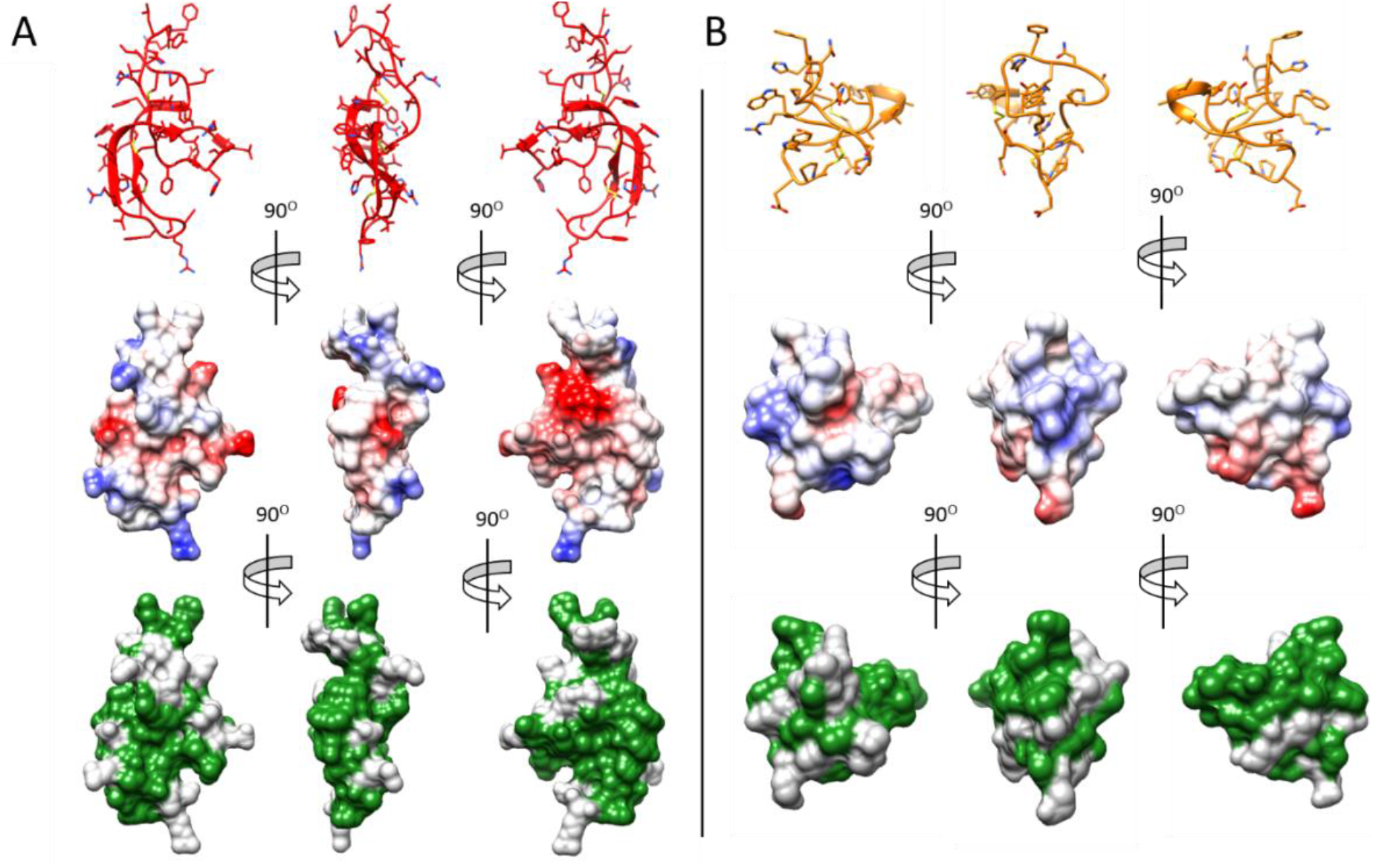
The hydrophobic paratopes of knob domains. The K8 and K92 knob domain peptides are shown in panels a and b, respectively. For each, the binding face is shown in the first column, with a 90° rotation shown in the second column and a further 90° rotation in the third columns shows the opposing, non-binding face of each domain. In the top row the amino acid side chains and disulphide bonds are shown in addition to the secondary structure. Using UCSF Chimera(Pettersen *et al.*, 2004), surface visualisation and coulombic colouring has been applied to denote charge in the middle row, with blue denoting positively charged residues and red negatively charged. The bottom row shows colouring of the non-polar residues, highlighting the hydrophobic nature of the knob domain paratopes.

### Crystal structure of the C5-K92 complex

We also present a crystal structure of the C5-K92 complex at a resolution of 2.75 Å (Supplementary 2.1 shows data collection and structure refinement statistics). Continuous electron density for the flexible C345C domain of C5 was observed, due to it being stabilised in an upward pose by crystal contacts, akin to the C5-RaCI-OmCI ternary complex structures (Protein Data Bank [PDB] accession codes: 5HCC, 5HCD and 5HCE(Jore *et al.*, 2016)). Despite being higher affinity than K8, density for K92 was not as well defined and so, to aid model building, we used mass spectrometry to perform a disulphide mapping experiment. The disulphide map of K92 identified formation of disulphide bonds between C9_K92_ and C23_K92_ and between C2_K92_ and C18_K92_ (Supplementary 2.8), enabling completion of the model. A mFo-DFc simulated annealing omit map of the C5-K92 complex is displayed in Supplementary 2.4, showing clear electron density for the peptide.

Similar to K8, K92 adopts a 3-strand β-sheet topology (Figure 2 c-d and Supplementary 2.9) but with only two disulphide bonds. With shorter β-strands and longer connecting loop regions, K92 exhibits a more compact, globular arrangement (Figure 3b). Two extended loop regions interact with C5, including an α-helix containing loop between β-strands 1 and 2, occupying a cleft between the MG1 and MG5 domains of the β-chain of C5. The 1365Å^2^ interaction surface is sustained via a sparse set of eight H-bonds (Supplementary 2.10). An elegant series of π-π and aliphatic–aromatic stacking interactions spans K92, encompassing: F26_K92_, H25_K92_, W21_K92_, W6_K92_ and P3_K92_, (Supplementary 2.11). From within this hydrophobic patch, important H-bonds occur between H25_K92_ and the backbone carbonyls of N77_C5_ and N81_C5_ on the MG1 domain.

The epitope for K92 is entirely contained within the binding interface of a previously reported immune evasion molecule, the 23 kDa SSL7 protein from *Staphylococcus aureus.* The C5-SSL7 crystal structure reveals the core of that interaction to be a series of five H-bonds between SSL7 and a region of β-sheet on the MG5 domain, spanning H511_C5_-E516_C5_ (Laursen *et al.*, 2010). Our structure shows that K92 also interacts with this area, inducing a re-orientation of the sidechain of H511_C5_ and forming a backbone H-bond with F510_C5_, but critically also makes interactions with the MG1 domain, effectively cross-linking the MG1 and MG5 domains. These changes beget different allosteric effects; SSL7, either in isolation or in complex with its second ligand IgA, is full, or occasional partial, antagonist of both the AP and CP(Bestebroer *et al.*, 2010; Laursen *et al.*, 2010), while K92 is a selective partial antagonist of the AP.

### The multi-purpose role of disulphide bonds

In the near absence of secondary structure, disulphide bonds are sources of stability for both peptides. For K92, both the backbone amide and carbonyl of C23_K92_ participate in H-bonds with the sidechain of S82_C5_, with close proximity to the electron dense disulphide bond lowering the conformational and desolvation entropy of these polar interactions. Within the knob domain paratopes, disulphide bonds also participate in sulphur-π interactions. For K8, an interchain sulphur-π stack between the C26_K8_-C40_K8_ disulphide bond and the aromatic of Y1378_C5_, elegantly positions the hydroxyl group of Y1378_C5_ to make a H-bond with D24_K8_. While for K92, an intra-chain sulphur-π stack between the C9_K92_-C23_K92_ disulphide bond and the aromatic of Y14_K92_ was used to orientate Y14_K92_, such that its hydroxyl group could participate in an interchain H-bond with N38_C5_.

Both K8 and K92 have prominent non-polar surface features (Figure 3). In addition to three disulphide bonds, K8 has several aromatic residues, F7_K8_, W9_K8_, Y11_K8_, F20_K8_, W26_K8_, Y29_K8_, Y38_K8_, F40_K8_, Y50_K8_ and F51_K8_, most of which occupy the opposing face and do not participate in the paratope. Hydrophobicity may be part of a specific strategy to maximise free enthalpy from the limited set of polar interactions. So, to evaluate bond enthalpy, we performed binding pose metadynamics(Clark *et al.*, 2016), an analysis typically employed to computationally evaluate the binding stability of chemical ligands(Fusani *et al.*, 2020). This *in silico* analysis suggested that both K8 and K92’s binding poses were exceptionally stable, with the interface maintaining the key interactions in spite of applied force (Supplementary 2.6 and 2.7). This, in conjunction with earlier kinetic studies(Macpherson *et al.*, 2020), highlights quality of the interactions made by both knob domains.

### Comparison to known antibody paratopes

Although antibody derived, K8 and K92 are structurally unique variable regions. We compared the K8 and K92 knob domains to a non-redundant set of 924 non-identical sequences of paired antibody-protein antigen structures from SAbDab(Dunbar *et al.*, 2014). Paratopes were defined as any antibody residues within 4.5 Å of the antigen in the structure. The paratopes of K8 and K92 contain 18 and 10 residues, respectively, which are within the typical range of antibody paratope sizes (Supplementary 2.14). Given this similarity in size, we searched for structurally and physicochemically similar antibody paratopes from the 924 antibody complexes but no similar paratope sites were found(Wong *et al.*, 2020). While the limited examples preclude firm conclusions, this lack of similarity could be due either to the unusual fold of the knob domains or to differences in paratope amino acid composition.

In terms of residue usage, one difference in paratope composition that is potentially universal is the presence of cysteine in the knob domains (Supplementary 2.15) which is uncommon in most antibody paratopes apart from broadly neutralising antibodies(Hutchinson *et al.*, 2019). Using Arpeggio(Jubb, 2015) to identify inter-(antigen contacting) and intra-paratope interactions revealed that on average, antibodies have 16 intra-paratope and 17 inter-paratope interactions; K8 is very close to this, with 15 intra-paratope and 17 inter-paratope interactions, whereas K92 paratope has fewer, with 9 intra-paratope and 10 inter-paratope interactions. A bovine Fab with an ul-CDRH3 was recently crystallised in complex with antigen, in this case a soluble portion of the HIV envelope(Stanfield *et al.*, 2020). While the low resolution of the crystal structure hindered analysis, a casual inspection of the paratope suggests 10 intra-paratope and 10 inter-paratope interactions are sustained by the knob domain, comparable to K92.

A search for structurally homologous proteins, using the DALI protein structure comparison server(Holm, 2020), did not find any 3-D structures similar to K8 or K92, including amongst the 14 structures of bovine Fabs with ul-CDRH3 in the PDB, potentially as a consequence of knob domains being shaped by their antigen.

### Solution techniques reveal allosteric networks

To contextualise our functional and structural data, we analysed the C5-knob domain complexes by two solution biophysical techniques – small angle X-ray scattering (SAXS) and hydrogen deuterium exchange mass spectrometry (HDX-MS).

SEC-SAXS, where size exclusion chromatography (SEC) immediately precedes the solution X-ray experiment ensuring a monodispersed sample, was performed in concert with SEC-MALLS (multi angle laser light scattering). Data were collected for C5 and the C5-K8, C5-K57, C5-K92 and C5-K149 complexes. (Figure 4a-c). SEC-MALLS confirmed that the increases in molecular weight of the complexes were consistent throughout the elution peaks (Supplementary 3.1 and 3.2). Interestingly, while SEC-SAXS elution profiles gave stable estimates of the radius of gyration (R_G_) across the tip of the peak, frames from the descending elution peaks show lower R_G_ values, suggesting either the presence of unbound C5 in the sample, which our SEC-MALLS data would preclude, or two distinguishable conformational states.

**Figure 4.**
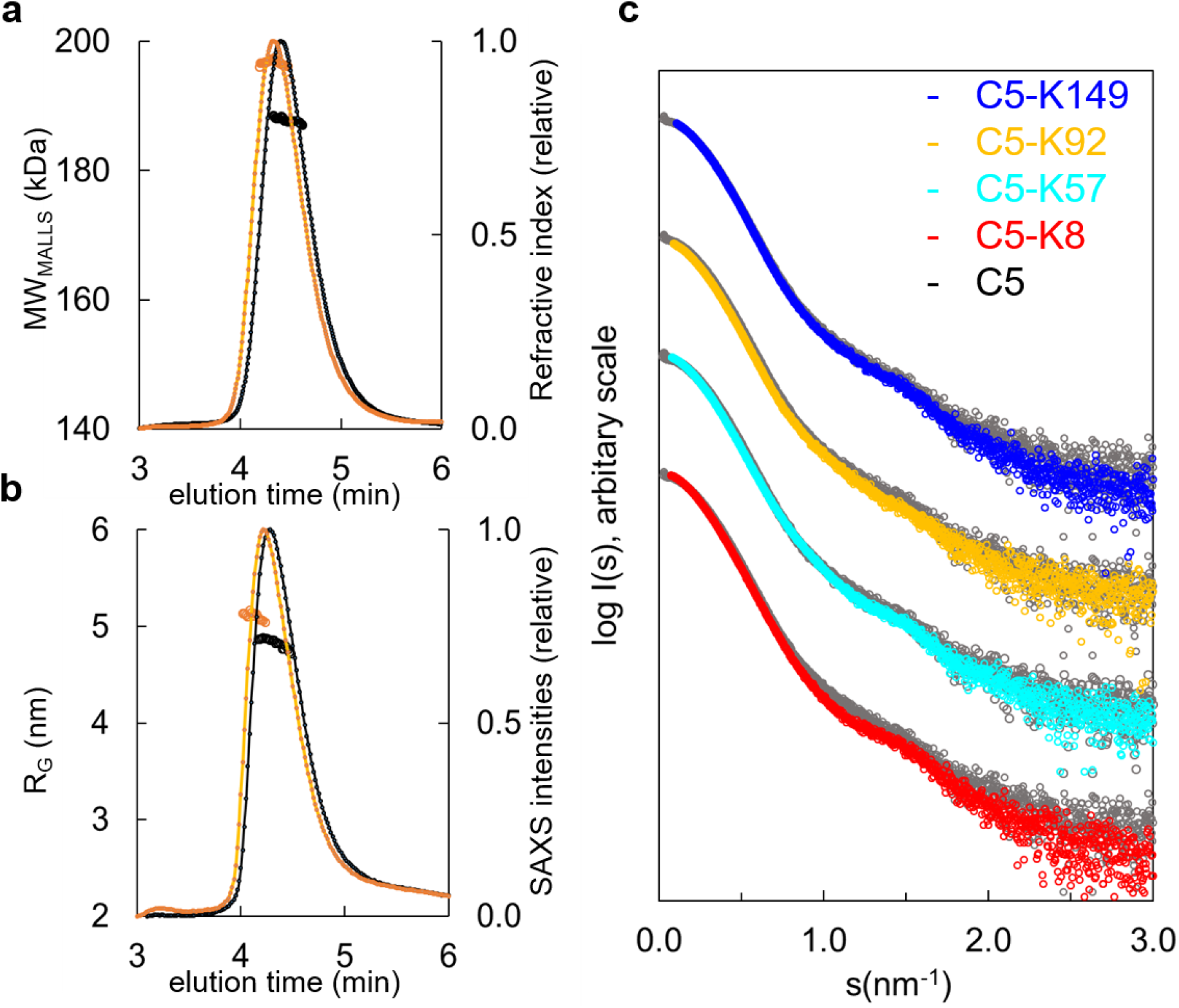
Hydrodynamic properties and solution conformation of C5 and C5-knob domain complexes by SAXS. SEC-MALLS chromatograms (a) for apo C5 (black) and C5-K92 (orange), show a homogenous molecular weight increase across the C5-K92 elution peak. The SEC-SAXS elution profile collected under identical experimental conditions (b) shows an increase in radius of gyration (R_G_) for the C5-K92 complex. Scattering curves of all C5-knob domains are shown (c), the C5-knob domain complexes are shown against apo C5 (in grey), for ease of viewing the curves are arbitrarily shifted in the Y axis.

Frames corresponding to the tip of the peak were averaged and submitted for full SAXS analysis. For the complexes, the scattering curves showed slight increases in both the RG and solute volume (Supplementary 3.1), with the C5-K8 complex showing the largest change and C5-K149 the smallest change, corresponding with the absence of function and suggesting K149 binds peripherally to a conformation closely resembling apo C5. For K92 and K57, the discrepancies observed in the mid s range indicate an overall change in flexibility of C5 upon binding of these peptides, and this tuning of dynamics may contribute to their mechanism.

Consistent with earlier observations(Fredslund *et al.*, 2008), comparison of the apo C5 experimental data with the theoretical scattering curve revealed discrepancies in the lowest angle range, indicating C5 adopts a more elongated conformation in solution than the crystal structure would suggest (Supplementary 3.3). To better approximate C5 in solution we performed a normal mode model analysis (NMA) using SREFLEX(Panjkovich and Svergun, 2016) and found that elongation of the C5 model improved the χ^2^ from >13 to 1.55. The fit of the C5-K92 complex was also markedly improved by the NMA (Supplementary 3.3), whereby elongation and incorporation of the peptide improved the model from an initial χ^2^ of >20, to 2.5 (with an overall root-mean-square [RMS] of 3.8 in both cases).

When using the C5-K8 co-crystal structure for fitting of the C5-K8 SAXS data, the absence of the C345C domain was problematic. The generation of a hybrid model where the C345C domain was reinstated produced a poor fit (χ^2^ = 75). Despite significant remodelling of C345C, we were unable to describe the solution structure in more detail. As the data were collected in SEC mode, it is unlikely that increases in R_G_ values and volume are due to oligomerisation and instead may suggest increased flexibility around the C345C linker. The absence of clear electron density for the C345C domain in the crystal structure may be a consequence of K8 inducing additional flexibility to this region, which again could contribute to the efficacy of the peptide.

The discrepancies between the crystal structures and the solution scattering data indicate that while permitting elucidation of the molecular interaction of the epitopes, the constraints of the crystal lattice may impede the detection of more subtle, global changes, leading to underestimation of the conformational changes induced by the peptide.

To further explore such effects in solution, we used HDX-MS to provide molecular level information on local protein structure and dynamics. HDX-MS measures the exchange of backbone amide hydrogen to deuterium in the solvent, with the rate of HDX determined by solvent accessibility, protein flexibility, and hydrogen bonding. To interpret the impact of peptide binding on C5 structural dynamics, we performed differential HDX (ΔHDX) analysis, comparing C5-knob domain complexes to apo C5, where shielding of C5 residues through participation in a binding interface will prevent deuteration, while conformational changes may increase or decrease deuterium uptake, in relation to the degree of solvent exposure.

For C5-K8, the sole protected region of C5 corresponded to the epitope on the MG8 domain (L1380_C5_-E1387_C5_), although the interface was not entirely defined (Figure 5a). Additional conformational changes were observed in the neighbouring C5d domain which becomes more solvent exposed, suggesting K8 is affecting the dynamics of this domain.

**Figure 5.**
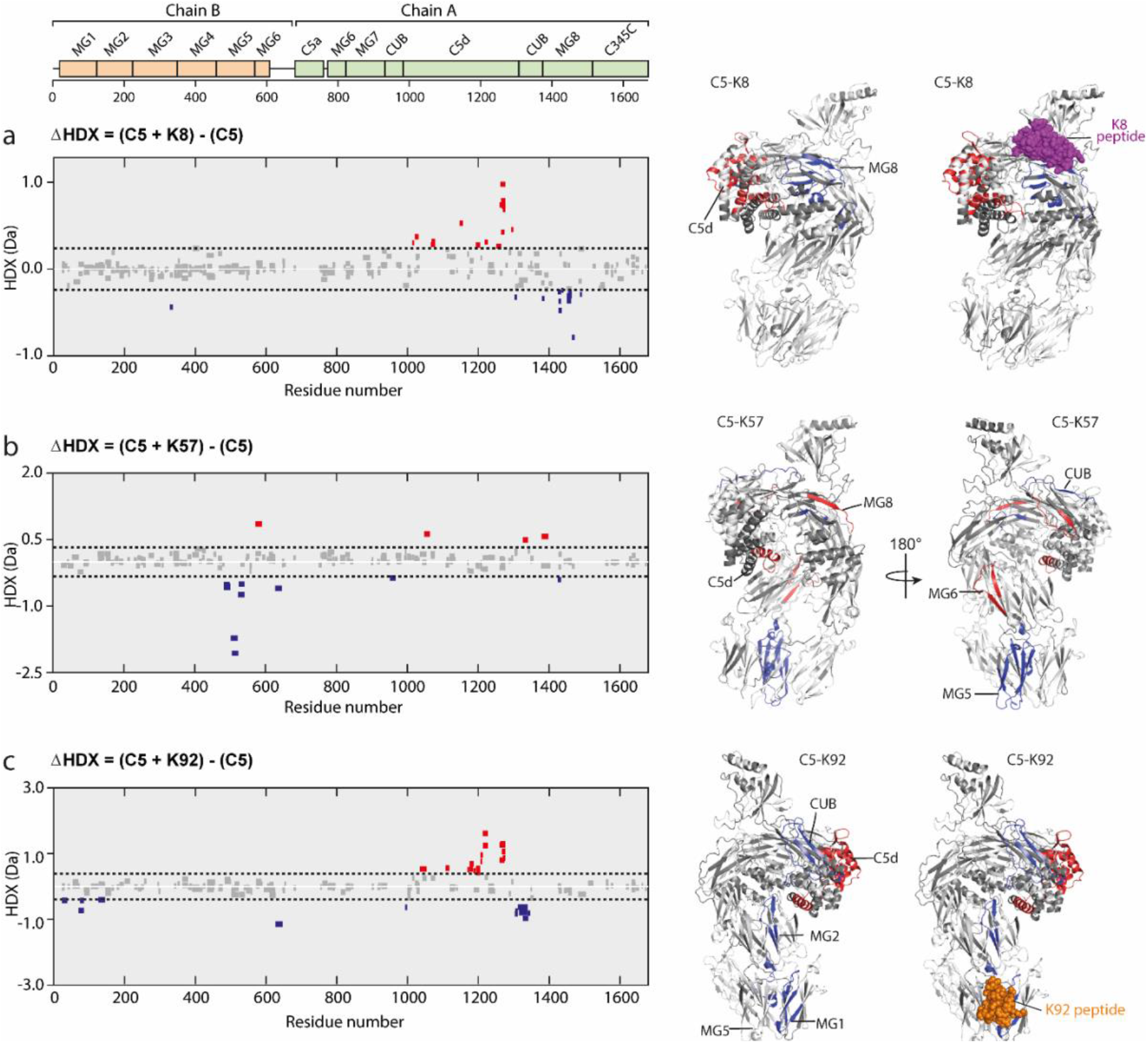
Impact of knob domain binding on the structural dynamics and conformation of C5. Differential HDX (ΔHDX) plots for C5 in complex with knob domains (a) K8, (b) K57 and (c) K92 at 1 hour of deterium exposure. Blue denotes peptides with decreased HDX (backbone H-bond stabilisation) and red denotes peptides with increased HDX (backbone H-bond destabilisation). 98% confidence intervals are shown as dotted lines. Peptides in grey have insignificant ΔHDX. Measurements were performed at in triplicate and all HDX-MS peptide data is detailed in Supporting Table XX. ΔHDX for C5 +K8, C5 + K57 and C5 + K92 are coloured onto C5 (PDB accession: 5HCC, minus OmCI and RaCI).

For the C5-K92 complex, consistent with the crystal structure, there was protection of the C5 residues located in the epitope between the MG1 and MG5 domains (H70_C5_-L85_C5_), shown in Figure 5c. There were also effects distal to the K92 binding site, notably in C5d (I1169_C5_-F1227_C5_) and neighbouring CUB domain (L1303_C5_-L1346_C5_), indicating a K92-induced conformational change. Interestingly, the allosteric network can be visualised by changes in solvent exposure which propagate from the K92 epitope through MG2 domain (L126_C5_-V145_C5_) and into the C5d and CUB domains. For the C5-K57 complex, the absence of a co-crystal structure meant we had no prior knowledge of the K57 epitope. However, clear protection was observed in the MG5 domain, immediately adjacent to the K92 epitope (N483_C5_-L540_C5_), with sparse areas of increased solvent exposure located in the MG6 (Q572_C5_-L590_C5_), MG8 (L1379_C5_-A1388_C5_) and C5d (K1048_C5_-Y1064_C5_) domains. A single protected peptide was also present in the CUB domain (G951_C5_-L967_C5_), suggesting the K57 epitope may be on either the MG5 or CUB domains.

The helical C5d domain is the target of two immune evasion molecules which have evolved in ticks, OmCI and RaCI, both of which inhibit C5 by crosslinking C5d to neighbouring domains(Jore *et al.*, 2016). Additionally, it has been shown that polyclonal antibodies raised against C5d inhibit binding of C5 to C3b(DiScipio, 1992). The binding site of OmCI is contained within the CUB and C5d domains, with only a single, non-bonded interaction to the C345C domain visible in the crystal structure(Jore *et al.*, 2016), which appears mediated by crystal contacts. This may suggest that K92 and K8 achieve efficacy in a manner similar to OmCI by modulating the C5d and CUB domains, but, in the case of K92, at a range of over 50 Å. Such remote effects are not unprecedented; allosteric structural changes can be propagated at over 150 Å in response to drug binding(Haselbach *et al.*, 2017). SSL7, which similarly binds the MG1-MG5 domains is also a partial antagonist, suggesting it too has an allosteric mechanism but not one which is selective for the AP.

There was little protection or deprotection of proteolytic fragments of the C5a domain in any of the complexes, we therefore propose that the knob domain peptides do not act by inducing conformational changes which shield the scissile arginine bond. The binding of K8, however, may induce some flexibility of C5a, as only sparse continuous electron density was observed for the N-terminal the linker extending from MG6 to this domain in the crystal structure. Taken in the context of the other changes, notably in the C5d and CUB domains, it is more probable that they affect more global changes in C5 which lower the affinity for C3b or the C5 convertases.

### Experimental validation of cooperativity

We hypothesised that these conformational changes may manifest as cooperativity between the different knob domain epitopes. To test this, we performed a SPR cross blocking-experiment where, using a Biacore 8K, we saturated a C5 coated sensor chip with two 20 μM injections of knob domain peptide before injecting a different peptide at 20 μM to assess its capacity to bind. Saturation of C5 with the non-functional K149 did not prevent subsequent binding of K8, K57 or K92 (Figure 6d), suggesting K149 does not share an epitope with the other ligands, nor does it significantly perturb C5 such that the other binding sites are affected.

**Figure 6.**
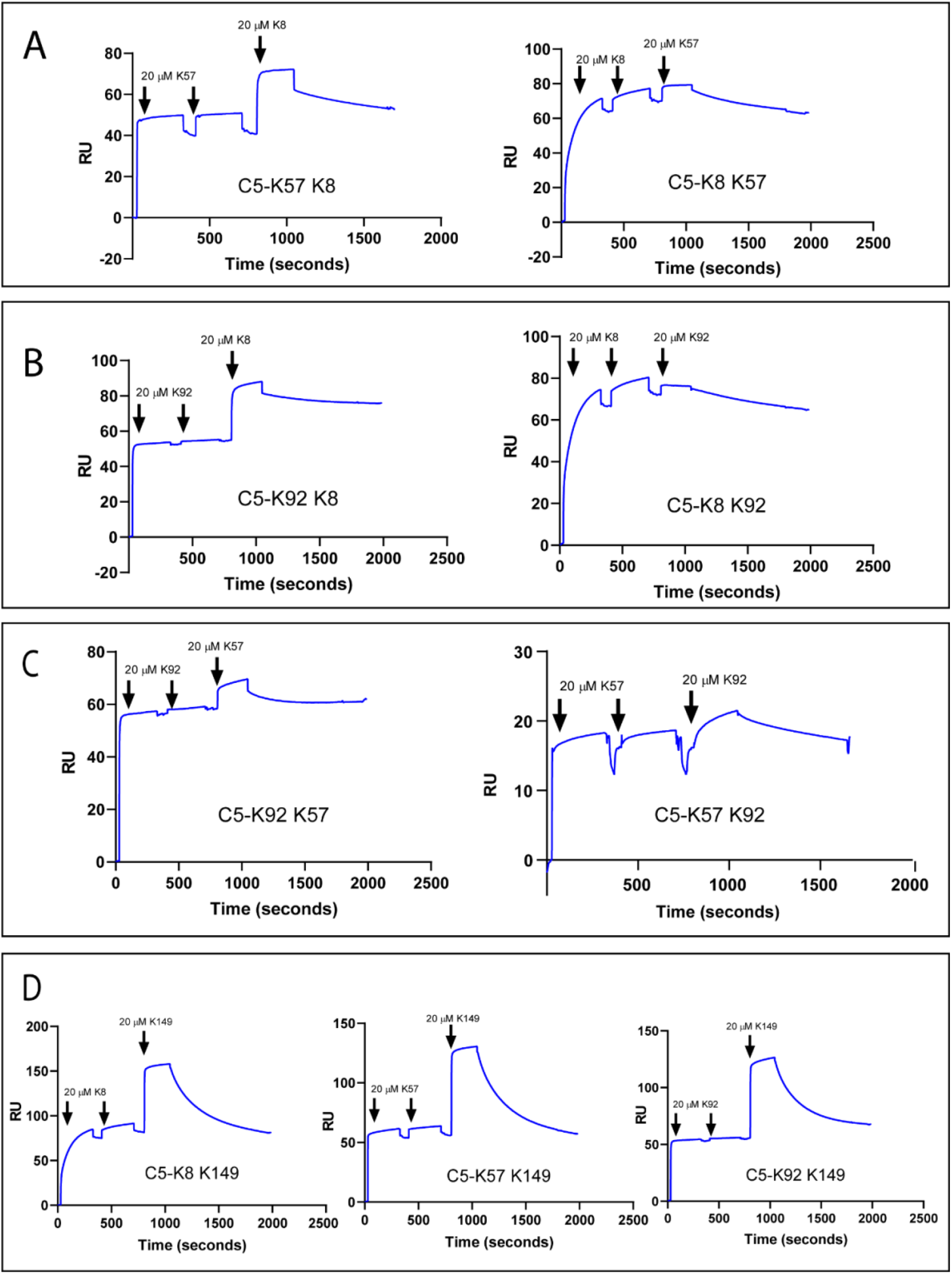
Surface plasmon resonance: cross blocking. Panels A, B and C highlight negative cooperativity between the K8, K92 and K57 peptides. Neither K57 or K92 can bind to the C5-K8 complex but K8 can bind, albeit at a lower level to, C5-K57 and C5-K92. We could not detect any negative cooperativity between K8, K57 or K92 with the silent binder K149, shown in panel D.

We detected negative cooperativity between the distal binding sites of K8 and K92, where, somewhat surprisingly, saturation of C5 with K8 entirely prevented binding of K92. Given the degree of separation between the K8 and K92 epitopes, in excess of 50 Å, and the conformational changes detected by HDX-MS analysis, we propose a non-steric mechanism whereby competition between K8 and K92 arises by negative cooperativity between the two distal epitopes. Correspondingly, when the order of addition was changed and C5 was saturated with K92, K8 was still able to bind, albeit to a lesser degree (Figure 6b). Saturation of C5 with K8 also entirely eliminated binding of K57, with a similar order of addition effect, whereby K8 could still partially bind to the C5-K57 complex (Figure 6a).

When C5 was saturated with K92 or K57 only very small amounts of subsequent binding of either peptide were observed by SPR (Figure 6c). Suggesting that the epitopes do not overlap but that considerable negative cooperativity exists. To further home in on the K57 binding site we measured binding to C5b in a SPR single-cycle kinetics experiment (Supplementary 5.1). Upon cleavage of C5a the remaining domains of the α-chain undergo a substantial conformational change, mediated by rearrangement of the MG8, CUB and C5d domains(Hadders *et al.*, 2012; Aleshin *et al.*, 2012). By SPR, K8 did not bind C5b, a likely consequence of conformational rearrangement of the CUB domain. However, K57, K92 and K149 all bound C5b with equal affinity to C5 (Supplementary 5.2). As the CUB domain is significantly altered in C5b, this increases the likelihood that, of the two protected regions identified by HDX-MS, the K57 epitope is on the MG5 domain.

By HDX-MS and SPR, we saw considerable long-range effects with all knob domains, with the exception of non-functional K149. We therefore suggest that knob domain peptides may be well suited to the allosteric modulation of proteins and that C5 may be particularly susceptible to modulation by such approaches.

## Discussion

We present knob domain peptides as novel therapeutic molecules, which rely on the bovine immune system to achieve high affinity binding through optimisation of amino acid composition, 3-D structure and disulphide bond network. Knob domains have only been recently isolated as a practicable antibody fragment(Macpherson *et al.*, 2020) and therefore structural information will greatly aid their development as therapeutics.

Due to the apparent structural homology of knob domains with certain venomous peptides, of which conotoxins and spider venoms are examples; it has been proposed that the knob domains of ul-CDRH3 might be similarly predisposed to target the concave epitopes of ion channels. Likewise, structural homology with defensin peptides has garnered hypotheses regarding an improved ability to bind viral capsid coats. Indeed, bovine antibodies with ul-CDRH3 have been raised against the viral capsid of HIV with exceptional efficiency, given the challenging nature of the antigen(Sok *et al.*, 2017; Stanfield *et al.*, 2020). However, the study presented here shows that, in the case of C5, concave epitopes are not the knob domain’s sole preserve. Notably, the MG8 domain epitope of K8 offers a planar pharmacophore and, while the K92 epitope is more undulating, casual inspection of the C5 structure reveals numerous deeper cavities available.

We note that the structural architecture of the knob domains varies for the epitope. Their immune derivation means that, unlike cysteine-rich peptides derived from other natural sources, such as venoms, the bovine immune system can be used to define specificity for any antigen. Comparative structural analysis suggests knob domain paratopes are differentiated from conventional antibodies, offering a different binding architecture to the ubiquitous VHH antibody fragment. While firm conclusions are hampered by limited examples, the number of interactions does not seem dissimilar from a mAb, which might be beneficial for tackling high affinity protein-protein interactions.

Knob domains offer hydrophobic paratopes, from which binding affinities broadly comparable to a conventional mAb can be obtained. The latent hydrophobicity is a likely consequence of germline priming for cysteine mutations which means the conserved IGDH8-2 D-gene segment, which encodes the knob domain and majority of the stalk, contains an astonishing 15 glycine and 17 tyrosine residues, each of which can be mutated to cysteine with a single base pair change. We propose that hydrophobicity may be a strategy to minimise desolvation entropy and thereby maximise the value of a H-bond network, enabling the most ligand-efficient, antibody-derived binding domain described to date.

Our structures demonstrate that the importance of the network of disulphide bonds goes beyond a stabilising role. An apparent paucity of secondary structure would suggest that stabilisation of the domain is indeed critical, but disulphide bonds are also a source of electron density for the paratope. We show that this electron density mediates sulphur-π interactions between the disulphide bond and nearby aromatic residues mediating intra- and inter-chain interactions. Finally, the disulphide bonds contribute even more hydrophobicity to the paratope which can improve the free enthalpy of local polar interactions. For example, the backbone amide and carbonyl of C23_K92_ form interchain H-bonds with C5, capitalising on the stability and proximal electron density of the disulphide bond to lower conformational and desolvation entropy.

Functional characterisation at the level of individual complement pathways identified K57 as a novel C5 inhibitor, which is a fully efficacious inhibitor of the terminal pathway in response to both CP and AP activation, and a potential therapeutic candidate for complement mediated disorders. Additionally, the discovery of K149 as a ‘silent binder’ of C5 may be of considerable value as a non-inhibitory reagent for the detection of native C5.

This study used X-ray crystallography in concert with solution biophysics methods to elaborate the mechanisms underpinning allosteric inhibition of proteins. With the caveat of small sample size, half of the knob domains exemplified in this study were allosteric. K92 achieved selective inhibition of the alternative pathway through a non-competitive mechanism. To our knowledge, this is the first reported example of complement pathway specific inhibition through C5 and the first experimental evidence reported for mechanistic differences between the AP and CP C5 convertases. This suggests an expanded therapeutic scope for C5, whereby tuning of the conformational ensemble with allosteric compounds can bias activation to leave certain complement pathways intact. Complete inhibition of the terminal pathway has been shown to increase the susceptibility of eculizumab patients to *Neisseria meningitidis* infections(McNamara *et al.*, 2017). Selective inhibition of C5-cleavage by the AP C5-convertase, and not the CP C5-convertase, may partially preserve serum bactericidal activity thereby lowering the risk of meningococcal disease.

While we cannot definitively explain why K92 was selective for the AP, through HDX-MS, we detected changes in the distal CUB and C5d domains. As the mechanism of K92 is non-competitive which precludes a steric mechanism, efficacy might be achieved through modulation of the CUB and C5d domains, in a manner similar to the immune evasion molecule OmCI. SAXS analysis also suggests that K8, K57 and K92 increase the flexibility of C5 and effects on dynamics may be a contributing factor in realising efficacy. This was most pronounced for K8 which appears to substantially affect the conformational sampling of C345C.

We also saw an unexpected degree of negative cooperativity between the different knob domains, suggesting that all the functional knob domains perturb the conformational state of C5 such that changes are induced in different regions. K57, with an E_max_ of 100 % for both pathways, was not demonstrably allosteric but still showed cooperativity with other functional knob domains, suggesting it stabilises a conformation of C5 that is less energetically favourable for binding of the other ligands. In HDX-MS experiments, K57 induces changes which span the length of C5, in a similar manner to K92, but the nature of the changes, and route of travel, are distinct to each ligand.

Our observations with K92 suggest that further work may be required to elucidate the mechanism of action of another binder of the MG1 and MG5 domains, SSL7. Given that SSL7 can be a partial inhibitor(Laursen *et al.*, 2010), even with co-binding of IgA, this precludes a steric mechanism and invites biophysical studies in solution. Additionally, another tick-derived inhibitor, Cirp-T, was also recently reported as predominantly binding to the MG4 domain, with an orthosteric mechanism of action attributed. However, we note that published data only showed an E_max_ of < 90 % in AP driven assays(Reichhardt *et al.*, 2020), indicating it is an allosteric C5 inhibitor for the AP, and potentially also the CP, C5 convertase, which may merit further investigation. This study highlights the importance of orthogonal biophysical techniques for the accurate interpretation of crystal structures. Taking a solely crystallographic approach, the constraints of the crystal lattice would have masked many of the distal effects that we observed.

This study is the first application of knob domain peptides and reveals an unexpectedly high incidence of allosteric modulators of complement C5. It is entirely possible that C5 could be particularly susceptible to allosteric modulation, or that by combining solution biophysics methods we may have observed effects that are typically missed. This study introduces knob domain peptides as a new peptide modality with unexplored therapeutic potential for the modulation of proteins and protein-protein interactions.

## Methods

### Complement proteins

Human C5 was affinity purified using an E141A, H164A OmCI column(Macpherson *et al.*, 2018). Briefly, human serum (TCS biosciences) was diluted 1:1 (v/v) with PBS, 20 mM EDTA and applied to a 5 mL Hi-Trap NHS column (GE Healthcare), which contained 20 mg of E141A H164A OmCI protein, at a rate of 1 mL/minute. The column was washed with 5x column volumes (CV) of PBS, C5 was then eluted using 2 M MgCl2 and immediately dialysed into PBS. C5b was prepared from human C5 by incubating C5 with CVF, factor B and factor D, at a 1:10 molar ratio, as previously described(Jore *et al.*, 2016). C5a was removed using a spin column with 30 kDa cut-off (Thermo Fisher).

### Knob domain peptide production

Knob domain peptides were expressed fused to the CDRH3 of the PGT-121 Fab, as previously described(Macpherson *et al.*, 2020). Plasmid DNA for each construct was amplified using QIAGEN Plasmid Plus Giga Kits. Expi293F cultures were transfected with Expifectamine 293 Transfection kits (Invitrogen), as per the manufacturer’s instructions. The cells were cultured for four days, and supernatants harvested by centrifuged at 4000 rpm for one hour. Harvested supernatants were applied to a Hi-Trap Nickel excel columns (GE Healthcare) using an Akta pure (GE Healthcare). Cell supernatants were loaded at 2.5 mL/minute, followed by a wash of 7x CV of PBS, 0.5 M NaCl. A second wash with 7x CV of Buffer A (0.5 M NaCl, 0.02 M Imidazole, PBS pH 7.3) was performed and samples were eluted by isocratic elution with 10x CV of Buffer B (0.5 M NaCl, 0.25 M Imidazole, PBS (pH 7.3). Post elution, the protein containing fractions were pooled and buffer exchanged into PBS, using dialysis cassettes (Thermo Fisher).

For isolation of the knob domain peptide, PGT-121 Fab-knob peptide fusion proteins were incubated with tobacco etch virus (TEV) protease, at a ratio of 100:1 *(w/w),* for a minimum of 2 hours at room temperature. Peptides were purified using a Waters UV-directed FractionLynx system with a Waters XBridge Protein BEH C4 OBD Prep Column (300 Å, 5 μm, 19 mm x 100 mm). An aqueous solvent of water, 0.1 % trifluoroacetic acid (TFA) and an organic solvent of 100 % MeCN was used. The column was run at 20 mL/min at 40 °C with a gradient of 5-50 % organic solvent, over 11 minutes. Fractions containing knob peptide were pooled and lyophilised using a Labconco Freezone freeze drier.

### Complement activation assays

For the C3 and C9 ELISAs, microtiter plates (MaxiSorp; Nunc) were incubated overnight at 4°C with 50 μL of a solution of in 75 mM sodium carbonate (pH 9.6) containing either: 2.5 μg/ml aggregated human IgG (Sigma-Aldrich) for CP, or 20 μg/ml zymosan (Sigma-Aldrich) for AP. As a negative control, wells were coated with 1 % (w/v) BSA/PBS. Microtiter plates were washed four times with 250 μL of wash buffer (50 mM Tris-HCl, 150 mM NaCl and 0.1 % Tween 20 (pH 8) between each step of the procedure. Wells were blocked using 250 μL of 1 % *(w/v)* BSA/PBS for 2 hours at room temperature. Normal human serum was diluted in either gelatin veronal buffer with calcium and magnesium (GVB^++^:0.1 % gelatin, 5 mM Veronal buffer, 145 mM NaCl, 0.025 % NaN_3_, 0.15 mM calcium chloride, 1 mM magnesium chloride, pH 7.3; for CP) or Mg-EGTA (2.5 mM veronal buffer [pH 7.3] containing 70 mM of NaCl, 140 mM of glucose, 0.1 % gelatin, 7 mM of MgCl2, and 10 mM of EGTA; for AP). Serum was used at a concentration of 1 % in CP or 5 % in AP and was mixed with serially diluted concentrations of peptides (16 μM – 15.6 nM) in GVB^++^ or Mg-EGTA buffer, and preincubated on ice for 30 minutes. Peptide-serum solutions were then incubated in the wells of microtiter plates for 35 minutes for CP assays (both C3b and C9 detection) or 35 min for AP (C3b) or 60 min for AP (C9), at 37 °C. Complement activation was assessed through detection of deposited complement activation factors using specific antibodies against C3b (rat anti-human C3d, Hycult; HM2198) and C9 (goat anti-human C9, CompTech; A226), at a 1:1000 dilution. Bound primary antibodies were detected with HRP-conjugated goat anti-rat (Abcam; ab97057) or rabbit anti-goat (Dako; P0449) secondary antibodies, at a 1:1000 dilution. Bound HRP-conjugated antibodies were detected using TMB One solution (Eco-TEK) with absorbance measured at 450 nm.

For the C5b ELISA, assays were run using the CP and AP Complement functional ELISA kits (SVAR). For sample preparation: serum was diluted as per the respective protocol for the CP and AP assays. Serial dilutions of peptides were prepared and allowed to incubate with serum for 15 minutes at room temperature, prior to plating.

For the C5a ELISA, assays were run using the Complement C5a Human ELISA Kit (Invitrogen). For sample preparation: at the end of the 37 °C incubation of the serum/peptide samples on the C5b ELISA assay plate, 50 μL of the diluted, activated serum was transferred to a C5a ELISA assay plate containing 50 μL/well of Assay Buffer. All subsequent experimental steps were performed as described in the protocol.

### Haemolysis assays

GVB^++^ or Mg EGTA buffers, which had been supplemented with 2.5% glucose *(w/v),* were used for the CP and AP assays, respectively. For the AP, 150 μL of rabbit erythrocytes (TCS Biosciences) were washed twice, by addition of 1 mL of buffer and centrifugation at 800 xg for 1 minute, and finally resuspended in 500 μL of buffer. For the CP, 150 μL sheep erythrocytes (TCS Biosciences) were washed twice with 1 mL of buffer and sensitised with a 1/1000 dilution of rabbit anti-sheep red blood cell stroma antibody (Sigma; S1389). After a 30 °C / 30 minutes incubation, with shaking, the cells were re-washed and resuspended with 500 μL of buffer. Serial dilutions of peptide were prepared in the respective buffers and normal human serum was added at 1% for the CP and 4.5% for the AP (corresponding to CH50 of the serum). 90 μL of peptide-serum mixtures were plated into a V-bottom 96-well microtiter plate (Corning) and 10 μL of erythrocytes were added. Plates were incubated for 30 minutes at 37 °C, with shaking. Finally, 50 μL of buffer was added, the plates centrifuged at 800 xg, and 80 μL of supernatant was transferred to an ELISA plate (Nunc) and absorbance measured at 405 nm.

### Crystallography and structure determination

6.1 mg/ml C5 (20 mM Tris-HCl, 75 mM NaCl, pH 7.35) was mixed at a 1:1 molar ratio with either the K8 or K92 peptides. Crystallisation trials were initiated by the vapor-diffusion method at 18 °C with a 1:1 mixture of mother liquor (*v/v*). C5-K8 crystals were grown in a mother liquor of 0.1 M ADA, 14 % ethanol (*v/v*), pH 6.0. For C5-K92 crystals, the mother liquor was 0.1 M bicine/Trizma (pH 8.5), 10 % *(w/v)* PEG 8000, 20 % (*v/v*) ethylene glycol, 30 mM sodium fluoride, 30 mM sodium bromide, 30 mM sodium iodide(Gorrec, 2009). Prior to flash freezing in liquid nitrogen, C5-K8 crystals were cryoprotected in mother liquor with 30 % MPD (*v/v*). C5-K92 crystals were frozen without additional cryoprotection.

Data were collected at the Diamond Light Source (Harwell, UK), on beamline I03, at a wavelength of 0.9762 Å. The C5-K8 structure was solved using the automated molecular replacement pipeline Balbes(Long *et al.*, 2008) using the apo C5 structure (PDB accession code: 3CU7), minus the C345c domain. The C5-K8 complex crystallised in space group P2_1_2_1_2_1_ with 1 molecule in the asymmetric unit. A backbone model of the K8 peptide was produced using ARP-wARP(Langer *et al.*, 2008) which informed manual model building in Coot(Emsley *et al.*, 2010), within the CCP4 suite(Winn *et al.*, 2011). The model was subjected to multiple rounds of refinement in Refmac(Murshudov, Vagin and Dodson, 1997) and Phenix(Adams *et al.*, 2010). The overall geometry in the final structure of the C5-K8 complex is good, with 97.2 % of residues in favoured regions of the Ramachandran plot and no outliers.

The C5-K92 complex crystallised in space group C2 with 1 molecule in the asymmetric unit. C5 was solved by molecular replacement with Phaser(McCoy *et al.*, 2007) using the C5-OmCI-RaCI structure (PDB accession code: 5HCC), with OmCI and RaCI removed. Manual building of the K92 peptide in Coot was greatly informed by mass spectroscopy disulphide mapping experiments. The model was subjected to multiple rounds of manual rebuilding in Coot and refinement in Phenix(Adams *et al.*, 2010). The overall geometry in the final structure of the C5-K92 complex is good, with 95.2 % of residues in favoured regions of the Ramachandran plot and no outliers. Structure factors and coordinates have been deposited in the PDB (PDB accession codes 7AD6 and 7AD7). Crystal trials were also performed with the C5-K57 and C5-K149 complexes, but the resulting crystals diffracted poorly.

### Disulphide mapping of K92 peptide

250 μL K92 peptide at 1 mg/mL was reduced with 6 μL of 0.5 M Dithiothreitol (Thermo Fisher) for 40 minutes at 37 °C and alkylated with addition of 18 μL of 2-Iodoacetamide (Thermo Fisher), at room temperature for 30 minutes. Overnight dialysis into assay buffer (7.5mM Tris-HCl, 1.5mM CaCl2, pH 7.9) was performed using 2 kDa slide-a-lyzer cassettes (Thermo Fisher). Chymotrypsin (sequencing grade, Roche Applied Sciences) was reconstituted to 1 μg/μL in assay buffer and 5 μL of reconstituted enzyme was added to 80 μL of sample. Once mixed, the sample was incubated at 37 °C for 1.5 hours before being quenched with 5 μL of 1 % TFA. Samples were diluted 1 in 10 and 5 μL was loaded onto the analytical column.

Liquid Chromatography electrospray ionisation mass spectrometry was acquired using an Ultimate 3000 UHPLC system (Thermo Fisher Scientific, San Jose, CA) coupled with a Q-Exactive Plus Orbitrap (Thermo Fisher Scientific). Separations were performed using gradient elution (A: 0.1 % Formic acid, B: 0.1 % Formic acid in acetonitrile) on an Acquity UPLC BEH C18 Column (130 Å, 1.7 μm, 2.1 mm X 150 mm; Waters Corp., Milford, MA, USA) with the column temperature maintained at 40 °C.

The following analytical gradient at 0.2 mL/min was used: 1 % B was held for 2 minutes, 1-36 % B over 28 minutes, 36-50 % over 5 minutes, 50-99 % B over 0.5 minutes. There were sequential wash steps with changes in gradient of 99 %−1 % B over 0.5 minutes (at a higher flow rate of 0.5 mL/min) before equilibration at 1 % B for 6.5 minutes (at the original 0.2 mL/min).

A Full MS / dd-MS2 (Top 5) scan was run in positive mode. Full MS: scan range was 200 to 2000 m/z with 70,000 resolution (at 200 m/z) and a 3 × 10^6^ AGC target, 100 ms maximum Injection time. The dd-MS2: 2.0 m/z isolation window, CID fragmentation (NCE 28) with fixed first mass of 140.0 m/z, with a 17,500 resolution (at 200 m/z), 1 × 10^5^ AGC target, 200 ms maximum injection time. The source conditions of the MS were capillary voltage; 3 kV, S-lens RF level; 50, Sheath gas flow rate; 25, Auxiliary gas flow rate; 10, Auxiliary gas heater temp; 150 °C and the MS inlet capillary was maintained at 320 °C.

Data were acquired using XcaliburTM 4.0 software (Thermo Fisher Scientific), and raw files were analysed by peptide mapping analysis using Biopharma Finder 2.0 software (Thermo Fisher Scientific) by performing a disulphide bond search with a chymotrypsin (medium specificity) digest against the K92 peptide sequence. Assignments and integrations from Biopharma Finder were filtered to include only peptides identified as containing a single disulphide bond and with an experimental mass within |5| ppm of the theoretical mass. Intensities for all peptides containing the same cysteines pairing were summed and percentages were obtained from the summed against total intensities.

### Binding pose metadynamics

Simulation structures were prepared using Schrodinger’s Maestro Protein preparation wizard. The molecular dynamics runs were performed using the Schrodinger’s default implementation of the binding pose metadynamics with the peptide chain considered in place of a ligand. Additional RMSD calculations for the peptide internal structure assessment in the last 20% of the dynamics were performed relative to the starting structures.

### Small angle X-ray scattering (SAXS)

Data was collected at the EMBL P12 beam line (PETRA III, DESY Hamburg, Germany(Blanchet *et al.*, 2015)). Data was collected with inline size exclusion chromatography (SEC) mode, using the Agilent 1260 Infinity II Bio-inert LC. 50 μL of complement component C5 at 31.6 μM (5.96 mg/ml) was injected onto a Superdex 200 Increase 5/150 column (GE Healthcare) at a flow rate of 0.35 ml/min. The mobile phase was comprised of 20mM Tris pH 7.35, 75mM NaCl, and 3% glycerol. The column elute was directly streamed to the SAXS capillary cell, and throughout the 15-minute-run 900 frames of 1 sec exposure were collected. After data reduction and radial averaging the program CHROMIXS(Panjkovich and Svergun, 2018) was employed. Around 100 statistically similar buffer frames were selected and used for background subtraction of the sample frames from the chromatographic peak. This results in the final I(s) vs s scattering profiles, where s = 4πsinθ/λ, 2θ is the scattering angle and λ= 1.24 Å. The scattering data in the momentum transfer range 0.05 < s < 0.32 nm^−1^ were collected with a PILATUS 6M pixel detector at a distance of 3.1 m from the sample.

ATSAS 2.8(Franke *et al.*, 2017) was employed for further data analysis and modelling. The program PRIMUS(Konarev *et al.*, 2003) was used to perform Guinier analysis (lnI(s) versus s^2^) from which the radius of gyration, R_G_, was determined. Distance probability functions, p(r), were calculated using the inverse Fourier transformation method implemented in GNOM(Svergun, 1992) that provided the maximum particle dimension, D_max_. The concentration-independent molecular weight estimate, MWVC,is based on the volume of correlation(Rambo and Tainer, 2013). The values are reported in Supplementary 3.1.

Theoretical scattering profiles were computed from X-ray coordinates using Crysol(Svergun, Barberato and Koch, 1995) and SREFLEX(Panjkovich and Svergun, 2016) was used to refine the models. For this, the program partitions the structure into pseudo-domains and hierarchically employs NMA to find the domain rearrangements minimizing the discrepancy χ^2^ between the SAXS curve computed from the refined model and the experimental data.

On the same day, multi-angle laser light scattering (MALLS) data were collected with a separate SEC run under the same experimental conditions (set-up, buffer, run parameters etc.). For this, a Wyatt Technologies miniDAWN TREOS multiangle laser light scattering detector coupled to an OptiLab T-Rex differential refractometer for protein concentration determination (dn/dc was taken as 0.185) was used. The MALLS system was calibrated relative to the scattering from toluene. The MWMALLS distribution of species eluting from the SEC column were determined with the Wyatt ASTRA7 software package.

The experimental SAXS data and the models derived from them were deposited to the Small Angle Scattering Biological Data Bank (accession number: SASDJA6).

### Hydrogen/deuterium mass spectrometry

6 μM of C5 was incubated with 10 μM of peptide (K8, K92 or K57) to achieve complex during deuterium exchange conditions. 4 μL of C5 or the C5-peptide complex were diluted into 57 μL of 10 mM phosphate in H_2_O (pH 7.0), or into 10 mM phosphate in D_2_O (pD 7.0) at 25 °C. The deuterated samples were then incubated for 0.5, 2, 15 and 60 minutes at 25 °C. After the reaction, all samples were quenched by mixing at 1:1 (*v/v*) with a quench buffer (4 M Guanadine Hydrochloride, 250 mM Tris (2-carboxyethyl) phosphine hydrochloride (TCEP), 100 mM phosphate) at 1 °C. The final mixed solution was pH 2.5. The mixture was the immediately injected into the nanoAcquity HDX module (Waters Corp.) for peptic digest, using an enzymatic online digestion column (Waters Corp.) in 0.2 % formic acid in water at 20 °C and with a flow rate of 100 μL/min. All deuterated time points and un-deuterated controls were carried out in triplicate with blanks run between each data-point.

Peptide fragments were then trapped using an Acquity BEH C18 1.7 μM VANGUARD chilled pre-column for 3 min. Peptides were eluted into a chilled Acquity UPLC BEH C18 1.7 μm 1.0 × 100 using the following gradient: 0 minute, 5 % B; 6 minutes, 35% B; 7 minutes, 40 % B; 8 minutes, 95 % B, 11 minutes, 5 % B; 12 minutes, 95 % B; 13 minutes, 5 % B; 14 minutes, 9 5% B; 15 minutes, 5 % B (A: 0.2 % HCOOH in H2O, B: 0.2 % HCOOH in acetonitrile. The trap and UPLC columns were both maintained at 0°C. Peptide fragments were ionized by positive electrospray into a Synapt G2-Si mass spectrometer (Waters). Data acquisition was run in ToF-only mode over an m/z range of 50-2000 Th, using an MSE method (low collision energy, 4V; high collision energy: ramp from 18V to 40V). Glu-1-Fibrinopeptide B peptide was used for internal lock mass correction. To avoid significant peptide carry-over between runs, the on-line Enzymate pepsin column (Waters) was washed three times with pepsin wash (0.8 % formic acid, 1.5 M Gu-HCl, 4 % MeOH) and a blank run was performed between each sample run.

MSE data from undeuterated samples of C5 were used for sequence identification using the Waters Protein Lynx Global Server 2.5.1 (PLGS). Ion accounting files for the 3 control samples were combined into a peptide list imported into Dynamx v3.0 software (Waters). The output peptides were subjected to further filtering in DynamX. Filtering parameters used were: minimum and maximum peptide sequence length of 4 and 25, respectively, minimum intensity of 1000, minimum MS/MS products of 2, minimum products per amino acid of 0.2, and a maximum MH + error threshold of 10 ppm. DynamX was used to quantify the isotopic envelopes resulting from deuterium uptake for each peptide at each time-point. Furthermore, all the spectra were examined and checked visually to ensure correct assignment of m/z peaks and only peptides with a high signal to noise ratios were used for HDX-MS analysis.

Following manual filtration in Dynamx, confidence intervals for differential HDX-MS (ΔHDX) measurements of individual time point were calculated using Deuteros(Lau *et al.*, 2019) software. Only peptides which satisfied a ΔHDX confidence interval of 98 % were considered significant. The ΔHDX was then plotted onto the C5 structure in Pymol.

### Surface plasmon resonance, single cycle kinetics

On a Biacore 8K (GE Healthcare), human C5b was immobilised on a CM5 chip (GE Healthcare). Flow cells were activated using a standard immobilisation protocol: EDC/NHS was mixed at 1:1 ratio (flow rate, 10 μL/min; contact time, 30 seconds). C5b, at 1 μg/mL in pH 4.5 sodium-acetate buffer, was immobilized on flow cell two only (flow rate, 10 μL/min; contact time, 420 seconds). Finally, ethanolamine was applied to both flow cells (flow rate, 10 μL/min; contact time, 420 seconds). A final immobilization level of 500-700 response units was obtained.

Single-cycle kinetics were measured using a 7-point, 3-fold serial dilution (spanning a range of 1 μM to 1.4 nM) in HBS-EP buffer (GE healthcare). A high flow rate of 40 μL/min was used, with a contact time of 300 seconds and a dissociation time of 2700 seconds. Binding to the reference surface was subtracted, and the data were fitted to a single-site binding model using Biacore evaluation software. All sensorgrams were inspected for evidence of mass transport limitation using the flow rate-independent component of the mass transfer constant (tc).

### Surface plasmon resonance, cross-blocking

On a Biacore 8K (GE Healthcare), human C5, was amine coupled to a CM5 chip, using the same protocol as for C5b. A final immobilization level of approximately 1000-2000 response units was obtained. For cross blocking, the surface was saturated with two sequential injections of a 20 μM knob domain solution in HBS-EP buffer (GE healthcare), using a flow rate of 30 μL/min and contact time of 300 seconds. This was immediately followed with an injection of a second knob domain peptide, again at 20 μM in HBS-EP, with a flow rate of 30 μL/min and a contact time of 270 seconds, the dissociation phase was measured for 600 seconds. Binding to the reference surface was subtracted, and sensorgrams were plotted in Graphpad Prism software.

## Competing Interests

This work was performed by A.M. as partial fulfilment of the requirements for a PhD from the University of Bath and was funded by UCB. A.M., S.S., V.O., J.S., B.H., V.E., Z.A. and A.D.G.L are present or past employees of UCB and may hold shares and/or stock options. T.E.M is a Board member of Ra Pharmaceuticals, Inc. All other authors declare no competing interests.

## Acknowledgements

We would like to thank John Cashman for his help during crystallography screening.

## Author Contribution

A.M., A.D.G.L. and J.v.d.E. designed the study. The manuscript was written by A.M., A.D.G.L. and J.v.d.E., with input from all authors. A.M expressed and purified peptides. P.H.N. and T.E.M. purified complement proteins. In vitro assays were performed by A.M and M.L. Crystallography was performed by A.M with assistance from J.R.B, S.S, S.C and J.v.d.E. Disulphide mapping was performed by B.H. Comparative antibody analysis was performed by S.R., J.S. and C.M.D. Binding pose metadynamics was by V.O. SAXS experiments were performed by M.G. and D.S. and HDX-MS by Z.A. and V.E. SPR was performed by A.M. Supervision was by A.D.G.L and J.v.d.E.

## Supplementary Material

### Supplementary Section 1. Functional assays

| Construct | Geomean IC50 (nM) | Range (nM) | Average Emax (%) | Range (%) |
| --- | --- | --- | --- | --- |
| **K8<sup>a</sup>** | 9.3 | 3.0-24.5 | 68.2 | 59.5 – 79.56 |
| **K57** | 3.6 | 2.4 – 7.3 | 100.6 | 99.1 – 101.7 |
| **K92** | ND* | ND* | ND* | ND* |
| **K149** | ND* | ND* | ND* | ND* |

\*ND = Not detected.
<sup>a</sup> Data are an average from $n=6$

Alternative pathway C5b deposition ELISA. Data from $n=3$ , unless specified.
| Construct | Geomean IC50 (nM) | Range (nM) | Average Emax (%) | Range (%) |
| --- | --- | --- | --- | --- |
| **K8<sup>a</sup>** | 29.2 | 25.4 – 34.2 | 91.8 | 91.4 – 92.3 |
| **K57** | 30.4 | 22.7 – 52.1 | 110.3 | 97.9 – 134.0 |
| **K92<sup>b</sup>** | 32.4 | 31.5 - 33.1 | 62.14 | 58.0 - 67.7 |
| **K149** | ND* | ND* | ND* | ND* |

\*ND = Not detected.
<sup>a</sup> Data are an average from $n=5$
<sup>b</sup> Data are an average from $n=4$

Inhibition of classical pathway mediated C5a release. Data from $n=3$ independent titrations, unless otherwise stated
| Construct | Geomean IC50 (nM) | Range (nM) | Average Emax (%) | Range (%) |
| --- | --- | --- | --- | --- |
| **K8<sup>a</sup>** | 9.9 | 3.9 – 18.7 | 57.7 | 51.5 – 73.2 |
| **K57** | 4.0 | 2.8 - 5.6 | 98.7 | 95.9 - 100.6 |
| **K92** | ND* | ND* | ND* | ND* |
| **K149** | ND* | ND* | ND* | ND* |

\*ND = Not detected.
<sup>a</sup> Data are an average from $n=5$

Inhibition of alternative pathway mediated C5a release. Data from $n=3$ , unless otherwise stated
| Construct | Geomean IC50 (nM) | Range (nM) | Average Emax (%) | Range (%) |
| --- | --- | --- | --- | --- |
| **K8<sup>a</sup>** | 150.2 | 49.3 – 85.86 | 71.8 | 49.3 – 85.9 |
| **K57** | 26.6 | 25.0-27.7 | 97.8 | 95.1 - 101.3 |
| **K92** | 43.3 | 39.9 – 45.3 | 43.3 | 39.9 – 45.2 |
| **K149** | ND* | ND* | ND* | ND* |

\*ND = Not detected.
<sup>a</sup> Data are an average from $n=4$

**Inhibition of classical pathway mediated C9 deposition.**Data from $n=3$
| Construct | Geomean IC<sub>50</sub><br>(nM) | Range (nM) | Average Emax<br>(%) | Range (%) |
| --- | --- | --- | --- | --- |
| **K8<sup>#</sup>** | - | - | 63.4 | - |
| **K57** | 1.9 | 1.3-2.5 | 97.9 | 92.4 - 97.9 |
| **K92** | ND* | ND* | ND* | ND* |
| **K149** | ND* | ND* | ND* | ND* |

\*ND = Not detected.
<sup>#</sup> Hill slopes < 0.5 IC<sub>50</sub> values not reported.**Inhibition of alternative pathway mediated C9 deposition.**Data from $n=3$ , unless otherwise stated.

| Construct | Geomean IC<sub>50</sub><br>(nM) | Range (nM) | Average Emax<br>(%) | Range (%) |
| --- | --- | --- | --- | --- |
| **K8** | 232.4 | 131.3 – 521.5 | 73.3 | 61.2 – 90.74 |
| **K57** | 24.5 | 20.6 - 27.6 | 100.7 | 100.1 - 100.9 |
| **K92** | 791.9 | 759.8 - 846.7 | 44.2 | 40.9 – 45.9 |
| **K149** | ND* | ND* | ND* | ND* |

\*ND = Not detected.

**Inhibition of classical pathway haemolysis.**Data from $n=3$
| Construct | Geomean IC<sub>50</sub><br>(nM) | Range (nM) | Average Emax<br>(%) | Range (%) |
| --- | --- | --- | --- | --- |
| **K8<sup>#</sup>** | - | - | 95.6 | 91.7 – 99.7 |
| **K57** | 5.3 | 4.9 – 5.7 | 99.7 | 99.2 – 100.1 |
| **K92** | ND* | ND* | ND* | ND* |

\*ND = Not detected.
<sup>#</sup> curves not well described by a 4-PL fit, IC<sub>50</sub> values not reported.**Inhibition of alternative pathway haemolysis.**Data from $n=3$

| Construct | Geomean IC<sub>50</sub><br>(nM) | Range (nM) | Average Emax<br>(%) | Range (%) |
| --- | --- | --- | --- | --- |
| **K8** | ND* | ND* | ND* | ND* |
| **K57** | 402.7 | 218 – 660 | 102.2 | 90.9 – 109.6 |
| **K92** | 16.9 | 11.41 – 21.8 | 31.4 | 24.1 – 45.3 |

\*ND = Not detected.

### Supplementary Section 2. X-ray Crystallography

**Supplementary 2.1.** Table of data collection and structure refinement statistics

Supplementary Section 2. X-ray Crystallography Supplementary 2.1 Table of data collection and structure refinement statistics
|  | C5-K8 | C5-K92 |
| --- | --- | --- |
| **Data collection** |  |  |
| Space group | P 21 21 21 | C 1 2 1 |
| Cell dimensions |  |  |
| *a*, *b*, *c* (Å) | 81.56, 161.69, 187.38 | 204.27, 104.29, 154.88 |
| $\alpha$, $\beta$, $\gamma$ (°) | 90, 90, 90 | 90, 124.89, 90 |
| Resolution (Å) | 81.07-2.3 (2.382-2.3)<sup>a</sup> | 83.77-2.75 (2.848-2.75) |
| *R*<sub>merge</sub> | 0.043 (0.361) | 0.018 (0.204) |
| *I* / $\sigma I$ | 13.17 (2.07) | 28.79 (4.38) |
| CC<sub>1/2</sub> | 0.99 (0.67) | 0.99 (0.92) |
| Completeness (%) | 0.99 (0.99) | 0.99 (0.99) |
| Redundancy | 13.5 (13.0) | 13.9 (14.0) |
| **Refinement** |  |  |
| Resolution (Å) | 81.56-2.3 | 83.77-2.75 |
| No. reflections | 110479 (10914) | 69381 (6877) |
| *R*<sub>work</sub> / *R*<sub>free</sub> | 0.202/0.234 | 0.219/0.253 |
| No. atoms (non-H) | 12100 | 13095 |
| Protein | 11618 | 13024 |
| Ligand/ion | 80 | 59 |
| Water | 402 | 12 |
| *B* factors |  |  |
| Protein | 65.4 | 100.5 |
| Ligand/ion | 87.9 | 122.7 |
| Water | 51.9 | 65.3 |
| r.m.s. deviations |  |  |
| Bond lengths (Å) | 0.003 | 0.002 |
| Bond angles (°) | 0.62 | 0.58 |

<sup>a</sup>Values in parentheses are for highest-resolution shell.

**Supplementary 2.2.**
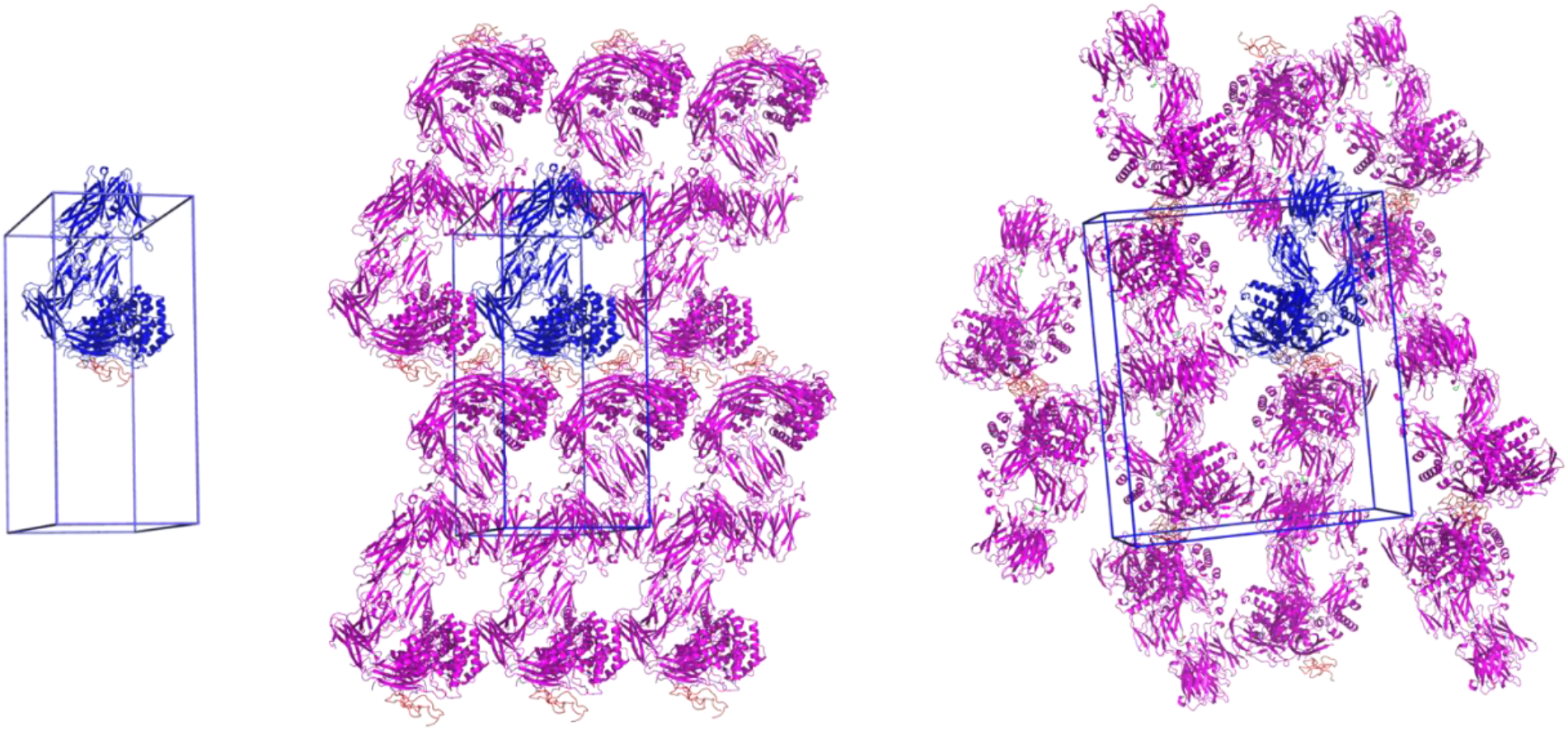
C5-K8 unit cell and crystal packing. A unit cell with a single molecule of the C5-K8 complex is shown followed by orthogonal views showing packing interactions as seen in the crystal lattice. C5-K8 is shown in cartoon representation with C5 coloured blue and K8 in orange. Symmetry-related molecules of C5 are coloured magenta with K8 in orange.

**Supplementary 2.3.**
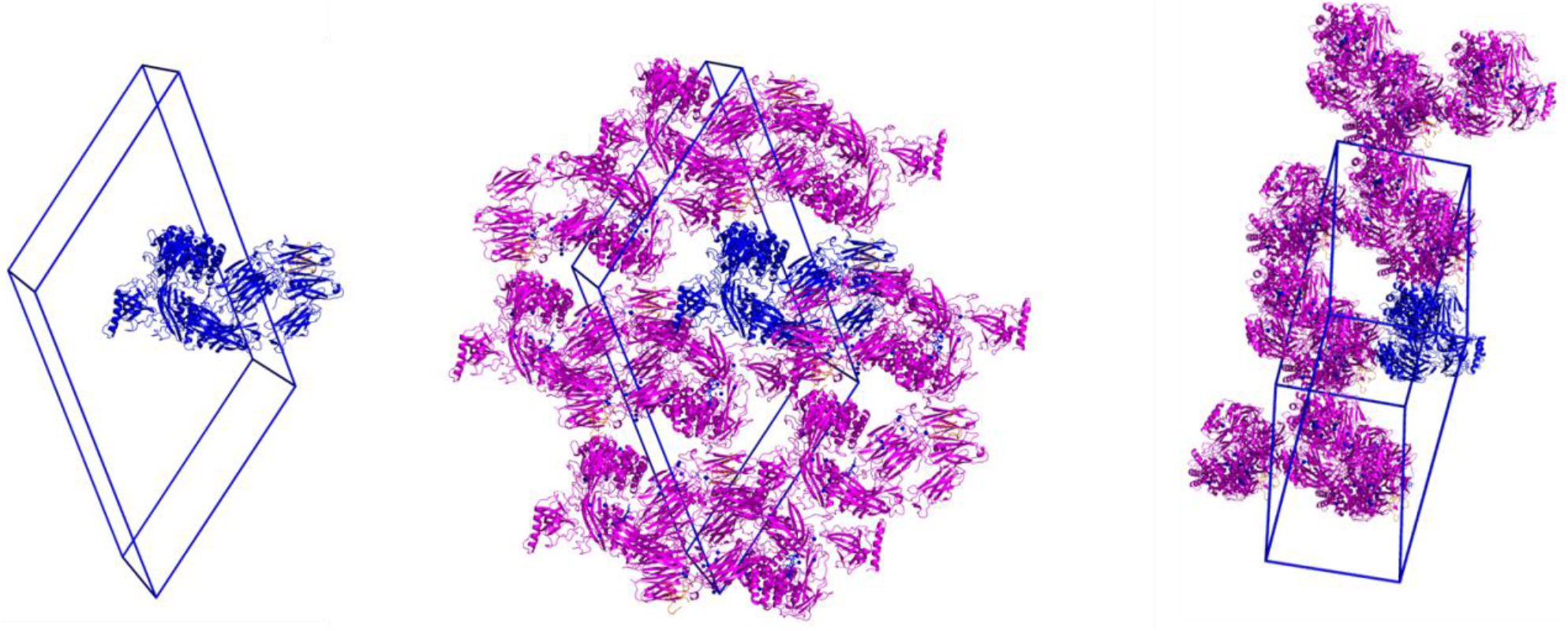
C5-K92 unit cell and crystal packing. A unit cell with a single molecule of the C5-K92 complex is shown followed by orthogonal views showing packing interactions as seen in the crystal lattice. C5-K92 is shown in cartoon representation with C5 coloured blue and K92 in orange. Symmetry-related molecules of C5 are coloured magenta with K92 in orange.

**Supplementary figure 2.4.**
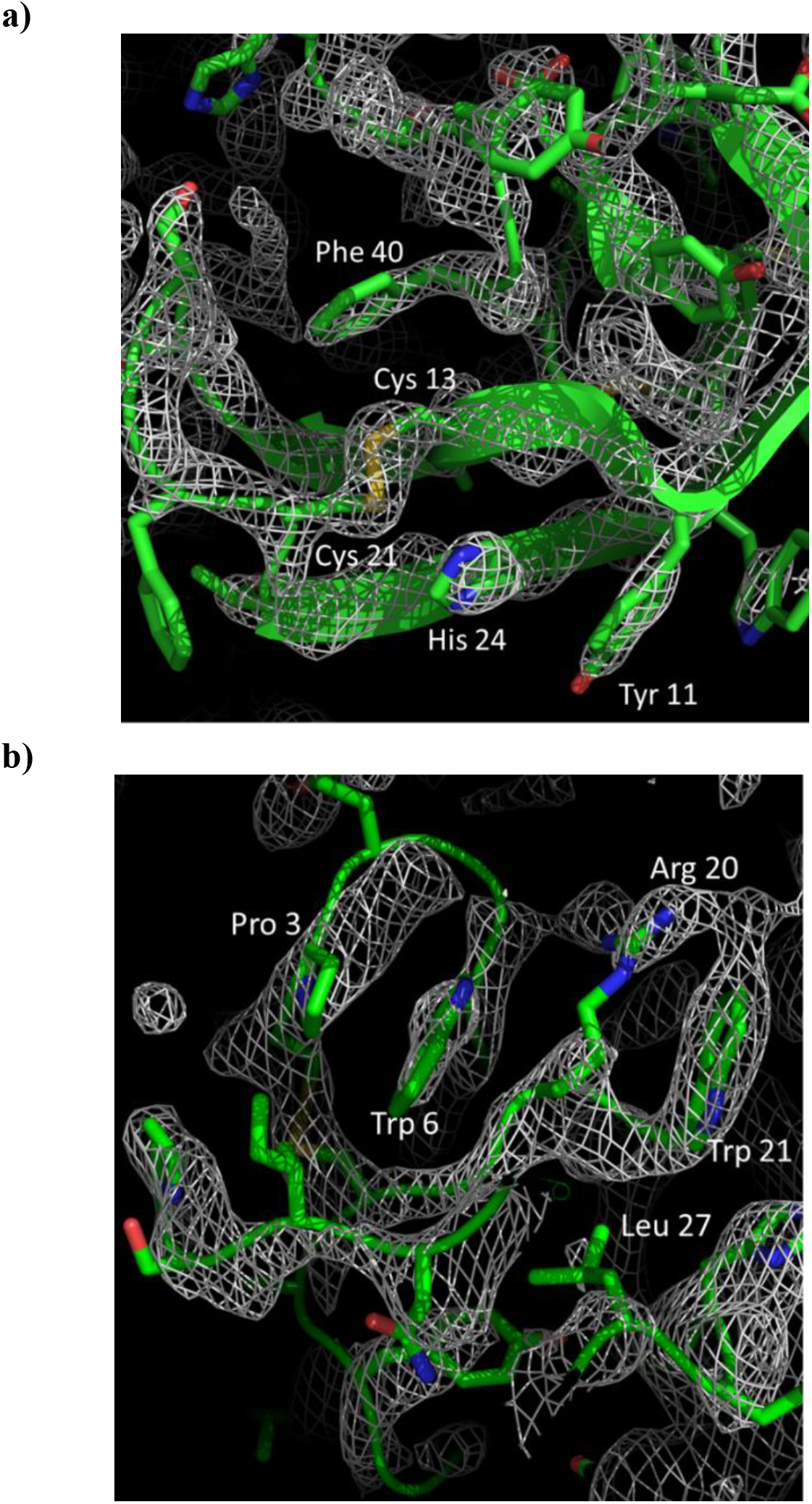
Simulated annealing omit maps of the C5-peptide complexes. The grey mesh shows mFo-DFc simulated annealing omit maps calculated in PHENIX and contoured at a) 3.0 sigma around the K8 peptide and b) at 1.0 sigma around the K92 peptide. In the omit calculation the peptides were deleted from the model. The peptides are displayed in green in cartoon representations with side chains shown as sticks and coloured according to the atom type (nitrogen in blue, oxygen in red, carbon in green and sulphur in yellow). Selected side chains are highlighted.

**Supplementary 2.5.**
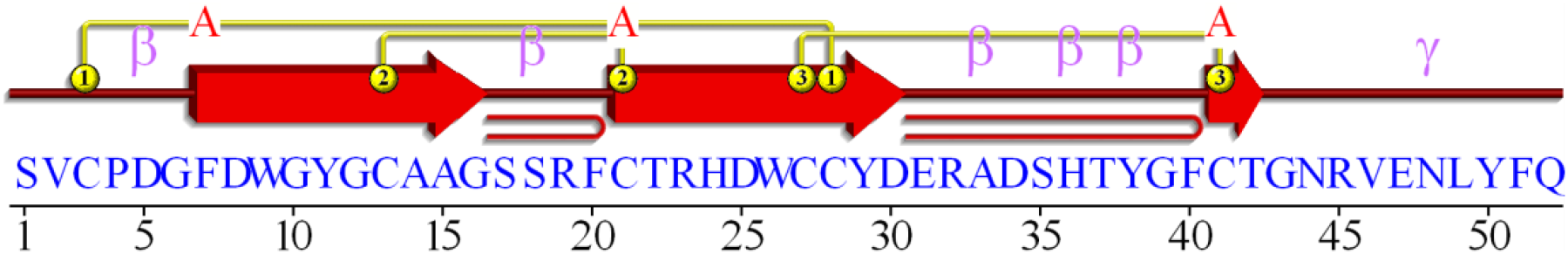
Structural topology of the K8 peptide.

**Supplementary 2.6.** Table of ionic interactions between K8 and C5. Ionic interactions as defined by the PDBePiSA macromolecular interfaces tool.

| K8 |  | C5 |  |
| --- | --- | --- | --- |
| Ionic interactions (5) |  |  |  |
| D25 | OD2 | NZ | K1409 |
| R32 | NH1 | OD2 | D1471 |
|  | NH2 | OD1/OD2 |  |
| H36 | NE2 | OD1 | D1382 |

**Supplementary 2.7.** Table of hydrogen bonds between K8 and C5. Hydrogen bonding interactions as defined by the PDBePiSA macromolecular interfaces tool.

| K8 |  | C5 |  |
| --- | --- | --- | --- |
| Hydrogen bonds (18) |  |  |  |
| A14 | O | N | L1379 |
| G16 | N | O | L1379 |
|  | O | N | I1381 |
| S18 | N | OD2 | D1382 |
|  | OG | NZ | K1380 |
| R23 | NH1 | O | E1373 |
|  |  | O | S1371 |
|  | NH2 | O | E1373 |
| D25 | OD2 | OH | Y1378 |
|  | OD2 | NZ | K1409 |
| R32 | NH1 | OD2 | D1471 |
|  | NH2 | OD1 |  |
| H36 | NE2 | OD1 | D1382 |
| G43 | N | OG | S1407 |
| N44 | N | O | S1469 |
|  | ND2 | O | S1470 |
| R45 | NH1 | O | S1411 |
|  | NH2 | O | E1414 |

**Supplementary 2.8.** Disulphide mapping of the K92 peptide. Table of intensities and resulting % (by total intensity) for the various peptides linked by a single disulphide bond as identified by Biopharma Finder.

| Cysteine Pairing | Intensity<sup>a</sup> | % (by total intensity) |
| --- | --- | --- |
| Cys23 to Cys9 | 150564035 | 62.71 |
| Cys18 to Cys2 | 85987873 | 35.81 |
| Cys23 to Cys2 | 2796033 | 1.16 |
| Cys23 to Cys23 | 527299 | 0.22 |
| Cys18 to Cys9 | 166812 | 0.07 |
| Cys23 to Cys18 | 17112 | 0.01 |
| Cys2 to Cys2 | 15354 | 0.01 |
| Cys18 to Cys18 | 11924 | 0.00 |
| Cys2 to Cys9 | 5193 | 0.00 |

<sup>a</sup> Sum of intensities of all peptides (containing a single disulphide bond) identified by Biopharma Finder as containing the same paired cysteines.

**Supplementary 2.9.**
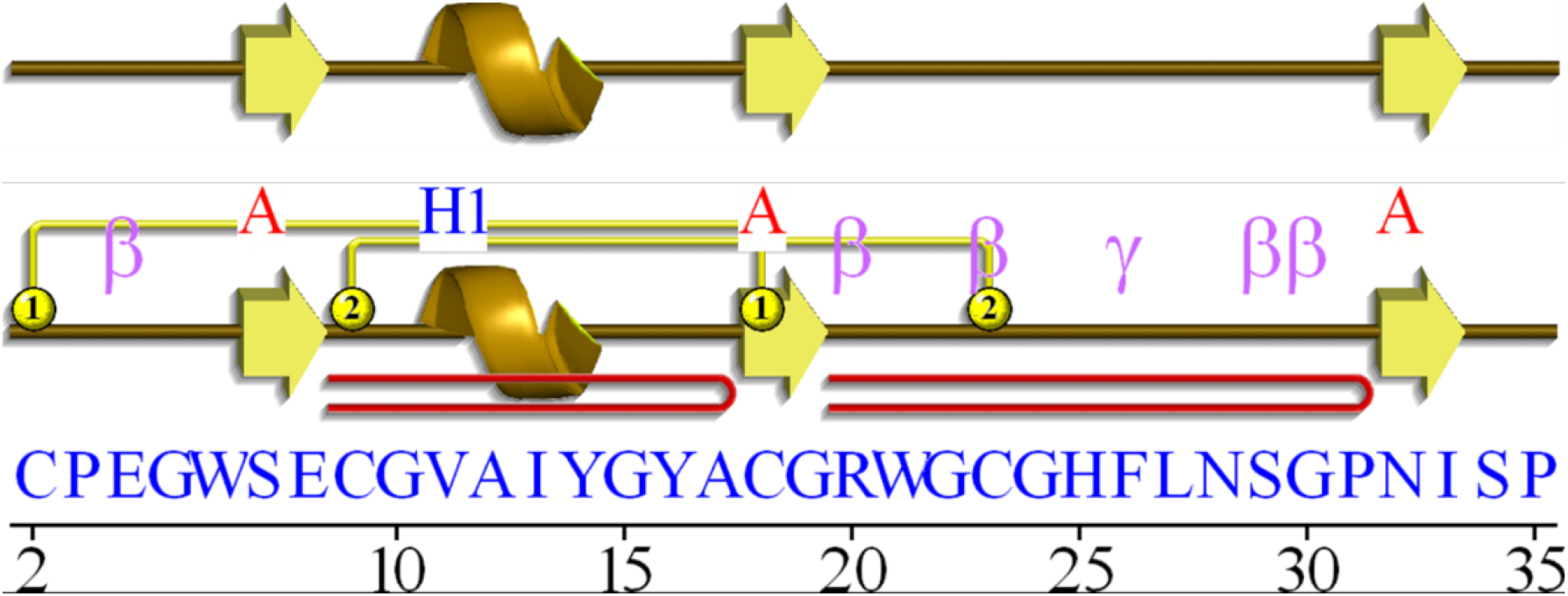
Structural topology of the K92 peptide.

**Supplementary 2.10.** Table of hydrogen bonds between K92 and C5. Hydrogen bonding interactions as defined by the PDBePiSA macromolecular interfaces tool.

| K92 |  | C5 |  |
| --- | --- | --- | --- |
| *Hydrogen bonds (8)* |  |  |  |
| G22 | N | OD1 | N81 |
| H25 | NE2 | O | A77 |
| F26 | N | O | F512 |
| I13 | O | ND2 | N533 |
| C23 | O | ND2 | N38 |
|  | O | OG | S82 |
|  | N | OG | S82 |
| G24 | O | N | F512 |

**Supplementary 2.11.**
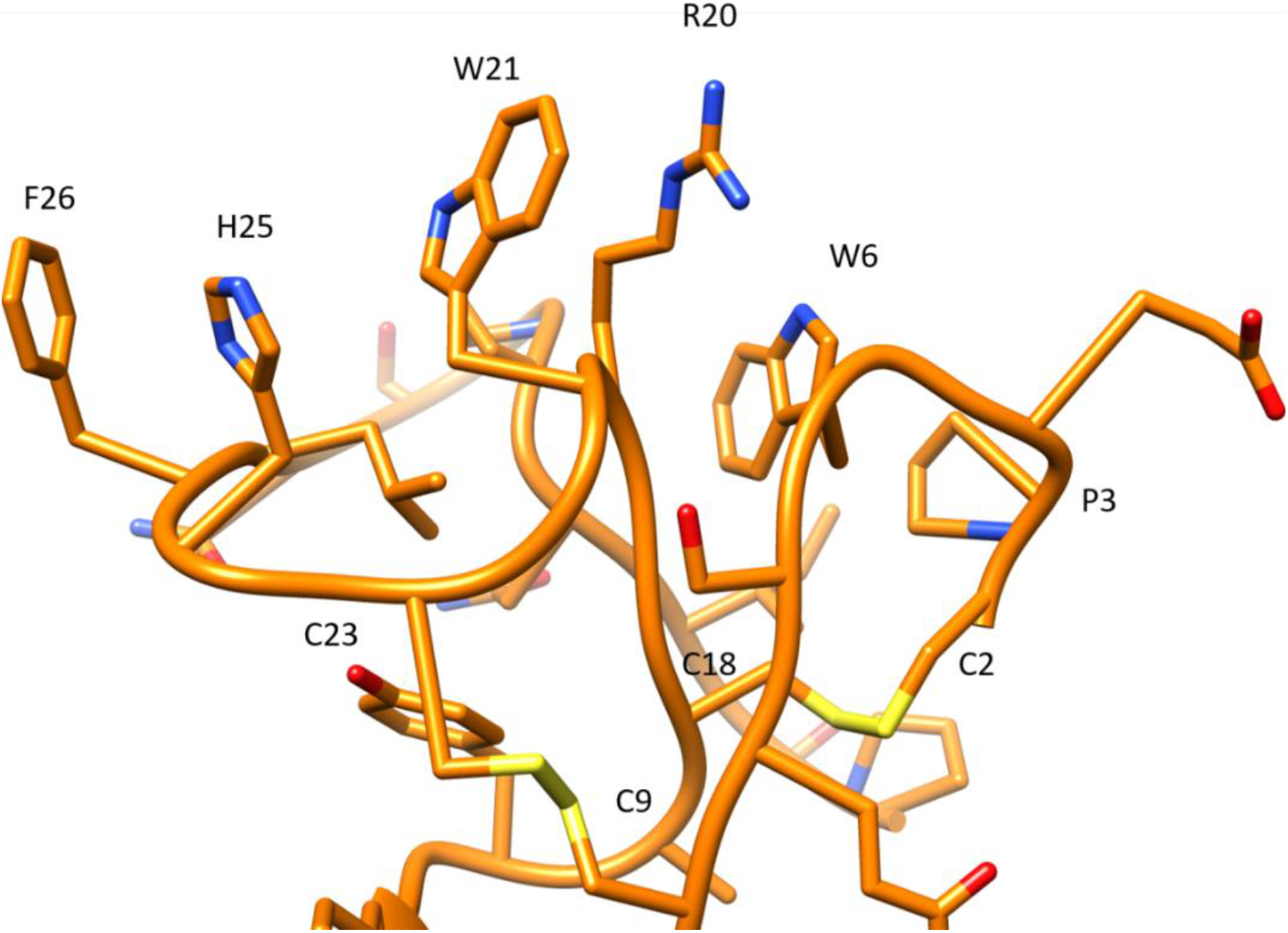
Stacking interactions of K92. An elegant series of π-π and π-aliphatic stacking interactions mark one face of the K92 peptide. The ordering of F26_K92_-H25_K92_-W21 _K92_-R20 _K92_-W6 _K92_-P3 _K92_ create a hydrophobic face through which decrease the desolvation entropy for hydrogen bonds made from the polar nitrogen of H25 _K92_ to C5.

**Supplementary 2.12.** Individual, total and average hydrogen bond persistence in a binding pose metadynamics simulation of K8-C5

| **HBond K8** | **HBond C5** | **Persistence score** |
| --- | --- | --- |
| 14:O | 1379:H | 1.000 |
| 16:H | 1379:O | 0.018 |
| 16O | 1381:H | 1.000 |
| 18:H | 1382:OD1(OD2) | 1.000 |
| 23:HH12(HH11,HH21,HH22) | 1373:O | 0.355 |
| 23:HH22(HH11,HH12,HH21) | 1373:O | 0.355 |
| 25:OD2(OD1) | 1378:HH | 0.964 |
| 32:HH12(HH11,HH21,HH22) | 1471:OD2(OD1) | 0.891 |
| 32:HH22(HH11,HH12,HH21) | 1471:OD2(OD1) | 0.891 |
| 43:H | 1407:OG | 0.609 |
| 44:HD22(HD21) | 1470:O | 0.900 |
| Total: |  | 7.98 |
| Average: |  | 0.726 |

**Supplementary 2.13.** Individual, total and average hydrogen bond persistence in a binding pose metadynamics simulation of K92-C5

| **HBond K92** | **HBond C5** | **Persistence score** |
| --- | --- | --- |
| 13:O | 533:HD21(HD22) | 0.909 |
| 22:H | 81:OD1 | 0.209 |
| 23:O | 38:HD22(HD21) | 0.536 |
| 24:O | 512:H | 0.536 |
| 25:HE2 | 80:O | 0.545 |
| Total: |  | 2.74 |
| Average: |  | 0.547 |

**Supplementary 2.14.**
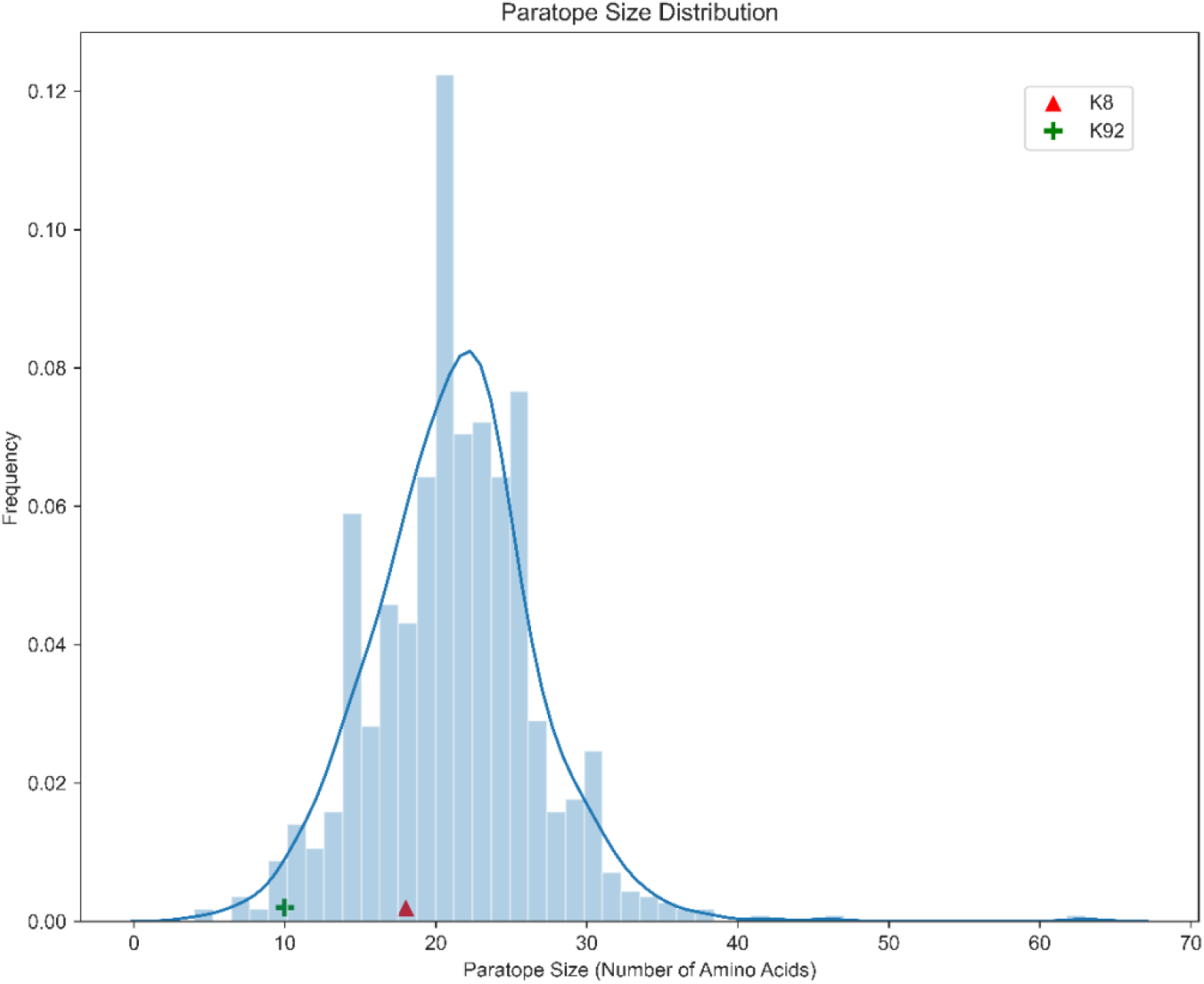
Paratope size. Comparison of K8 and K92 paratope size to nonredundant antibodies in SAbDab (N= 924). Paratopes were defined as antibody residues within 4.5Å of the antigen in the co-crystal structure.

**Supplementary 2.15.**
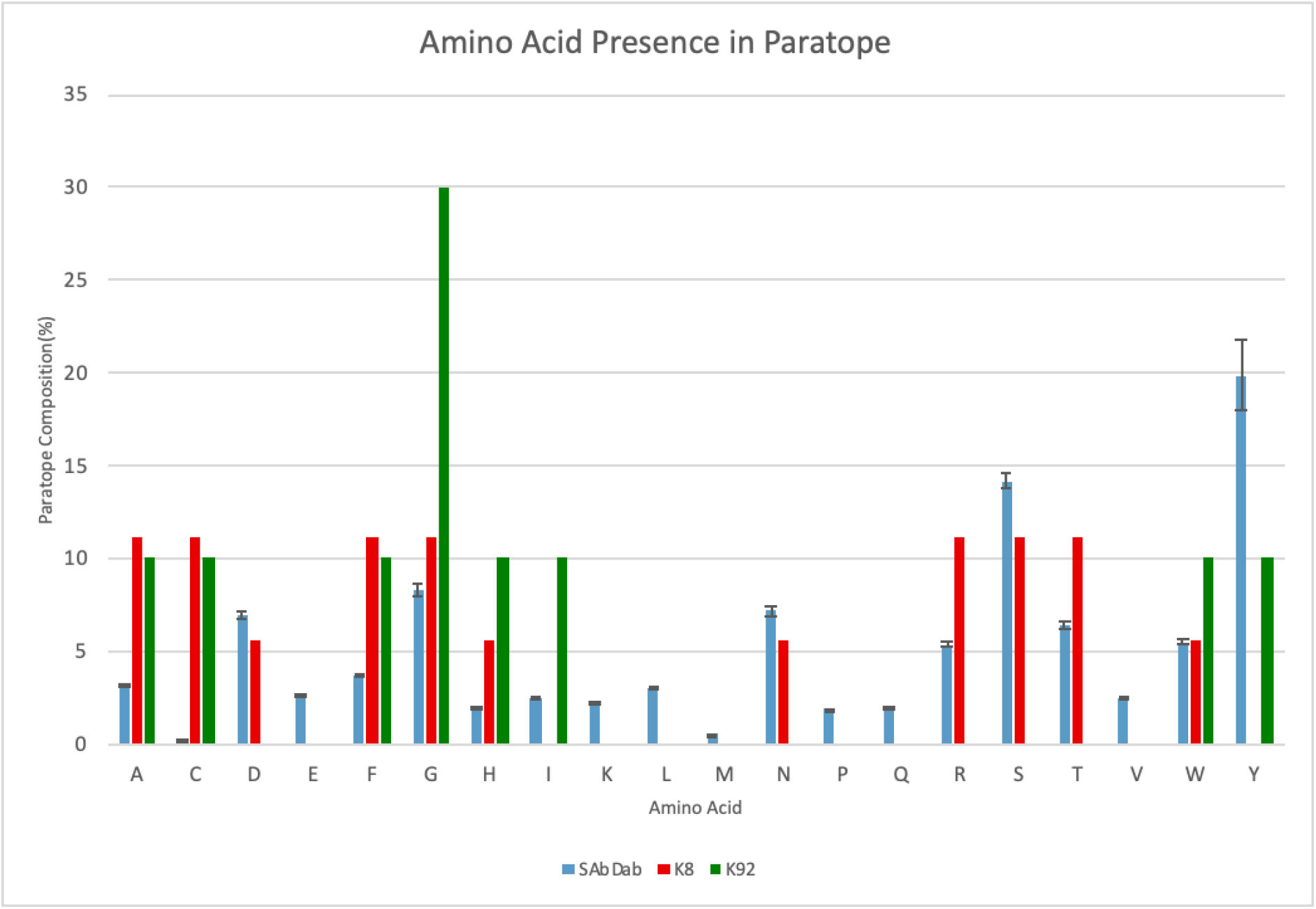
Amino acid presence in paratope. (defined as residues 4.5 Å from C5). Comparison of K8 and K92 paratope composition to nonredundant antibodies in SAbDab (n = 924). Paratopes were defined as antibody residues within 4.5Å of the antigen in the co-crystal structure. Error bars of 1x SD are included for SAbDab dataset.

### Supplementary Section 3. Small Angle X-ray Scattering

**Supplementary 3.1.** SAXS Summary Data Table

| Data collection parameters | C5 | C5-K8 | C5-K57 | C5-K92 | CK-149: |
| --- | --- | --- | --- | --- | --- |
| Instrument | EMBL P12 (PETRA III, DESY, Hamburg) |  |  |  |  |
| Beam geometry (mm<sup>2</sup>) | 0.2×0.12 |  |  |  |  |
| Wavelength (nm) | 12.4 |  |  |  |  |
| *s* range (nm<sup>-1</sup>) | 0.03-7.1 |  |  |  |  |
| Temperature (K) | 293 |  |  |  |  |
| Concentration range (ml) | 5.96 |  |  |  |  |
| SEC parameters |  |  |  |  |  |
| Column | Superdex 200 Increase 5/150 column |  |  |  |  |
| Buffer | 20mM Tris pH 7.35, 75mM NaCl, and 3% glycerol |  |  |  |  |
| Flow rate, duration | 0.35 ml/min; 15 min (900 frames) |  |  |  |  |
| Structural parameters |  |  |  |  |  |
| $R_g$ (nm) (from $P(r)$) | 4.8 ± 0.1 | 5.4 ± 0.1 | 5.2 ± 0.1 | 5.0 ± 0.1 | 4.9 ± 0.1 |
| $R_g$ (nm) (from Guinier plot) | 4.8 ± 0.1 | 5.4 ± 0.1 | 5.3 ± 0.1 | 5.1 ± 0.1 | 5.0 ± 0.1 |
| $D_{max}$ (nm) | 17.6 ± 0.5 | 18.3 ± 10 | 19 ± 0.5 | 19 ± 0.5 | 18.5 ± 0.5 |
| Porod volume estimate, $V_p$ (nm<sup>3</sup>) | 390 ± 10 | 450 ± 10 | 410 ± 10 | 410 ± 10 | 410 ± 10 |
| Molecular weight determination (kDa) |  |  |  |  |  |
| From volume of correlation, VC | 210 ± 20 | 240 ± 25 | 230 ± 25 | 220 ± 22 | 220 ± 22 |
| From *MALLS* | 188 ± 9 | 200 ± 10 | 199 ± 10 | 197 ± 10 | 195 ± 10 |
| Calculated monomeric *MW* from sequence | 186 | 192 | 190 | 191.5 | 190 |
| Software employed |  |  |  |  |  |
| Primary data reduction | SASFLOW |  |  |  |  |
| Data processing | CHROMIXS/PRIMUS |  |  |  |  |
| Computation of model intensities | CRY SOL |  |  |  |  |
| normal mode analysis | SREFLEX |  |  |  |  |

**Supplementary 3.2.**
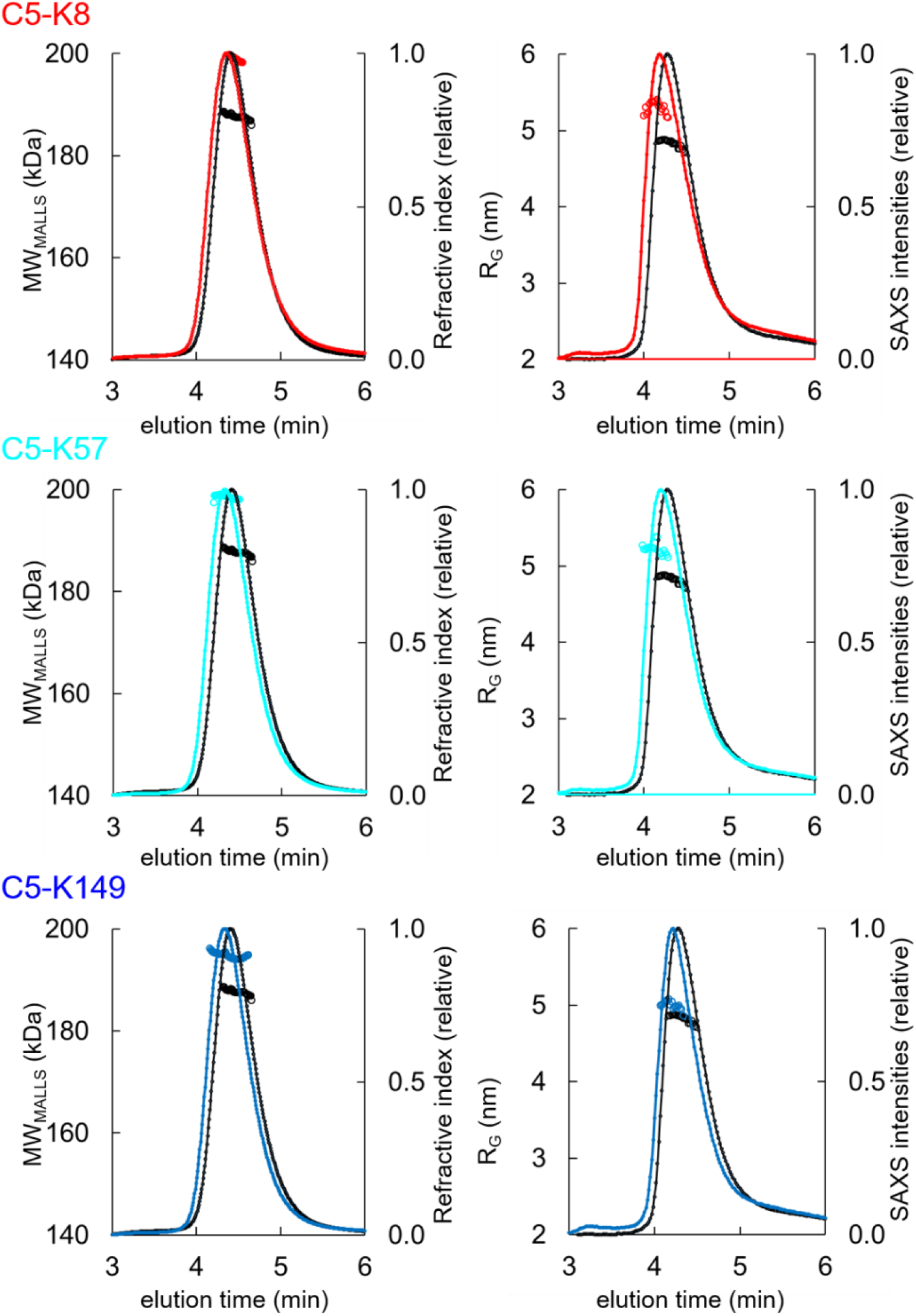
SEC-MALLS and SEC-SAXS chromatograms for C5-knob domain complexes. SEC-MALLS chromatograms for C5-knob domain complexes show a homogenous molecular weight increase across the elution peak. The SEC-SAXS elution profile collected under identical experimental conditions shows an increase in radius of gyration (R_G_) for the complexes.

**Supplementary 3.3.**
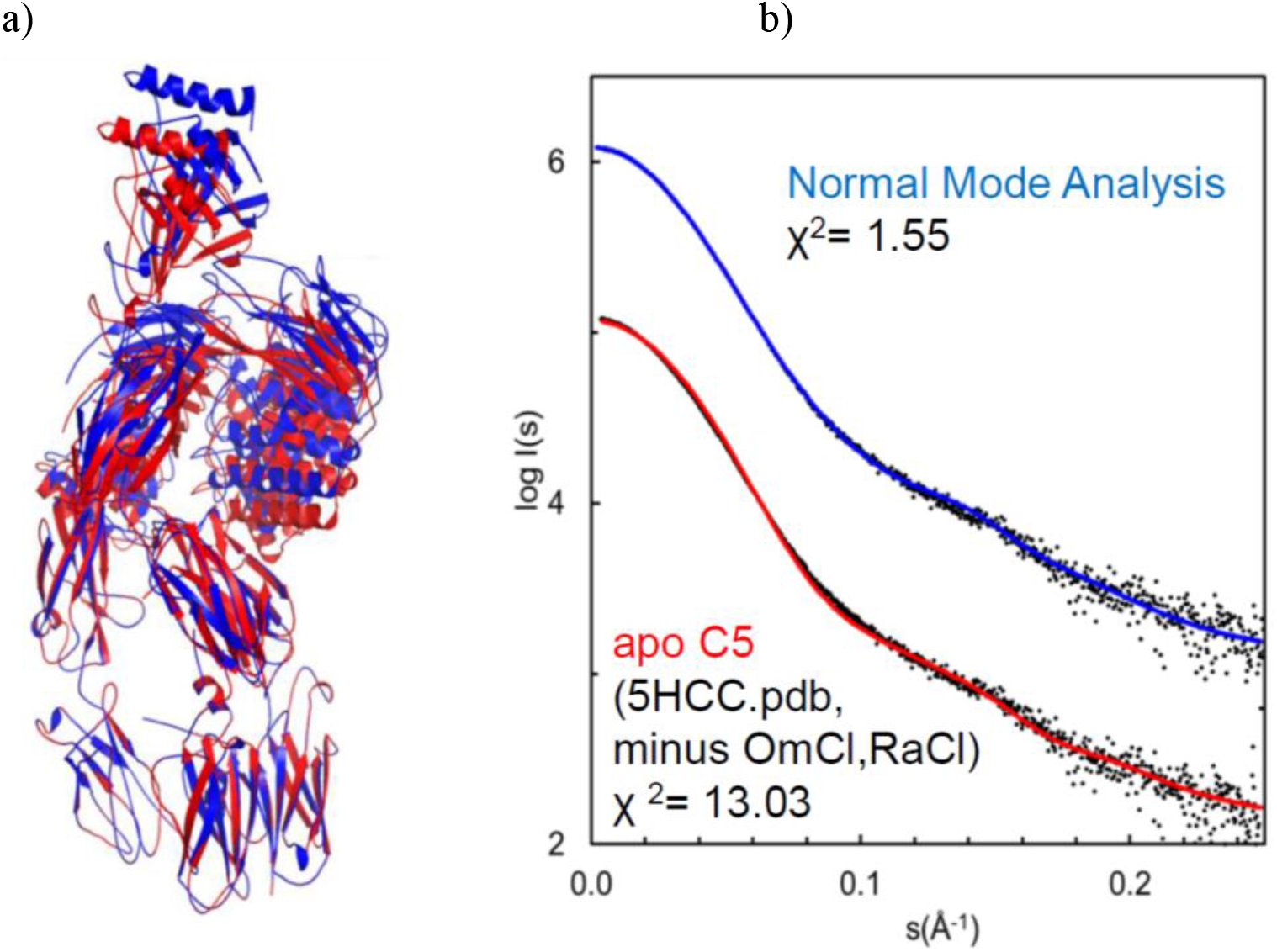
Solution structure of complement C5 in apo state. a) Overlay of cartoon representations of the C5 (red) and the model obtained with SREFLEX (normal mode analysis, blue). b) Right, the respective fits of the theoretical scattering curves to the SAXS data. χ^2^ values are indicated.

### Supplementary Section 4. HDX-MS

**Supplementary Figure 4.1.**
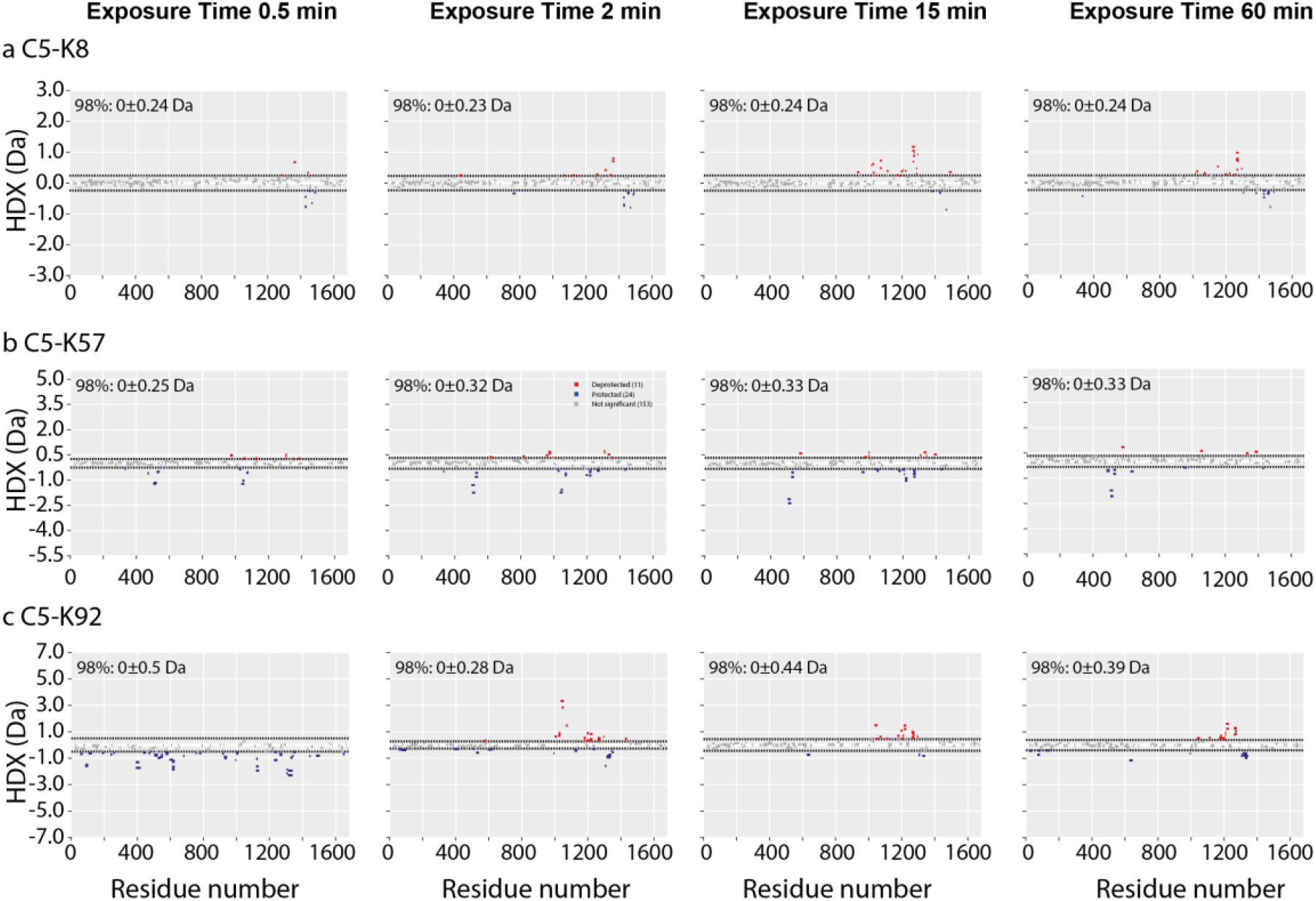
Woods plot displaying the differential HDX (ΔHDX) for C5 in complex with knob domains. (a) K8, (b) K57 and (c) K92 at four deterium exposure time points: 0.5 min, 2 min, 15 min and 1 hr. Blue denotes peptides with decreased HDX and red denotes peptides with increased HDX. 98% confidence intervals are shown as dotted lines.

**Supplementary Figure 4.2.**
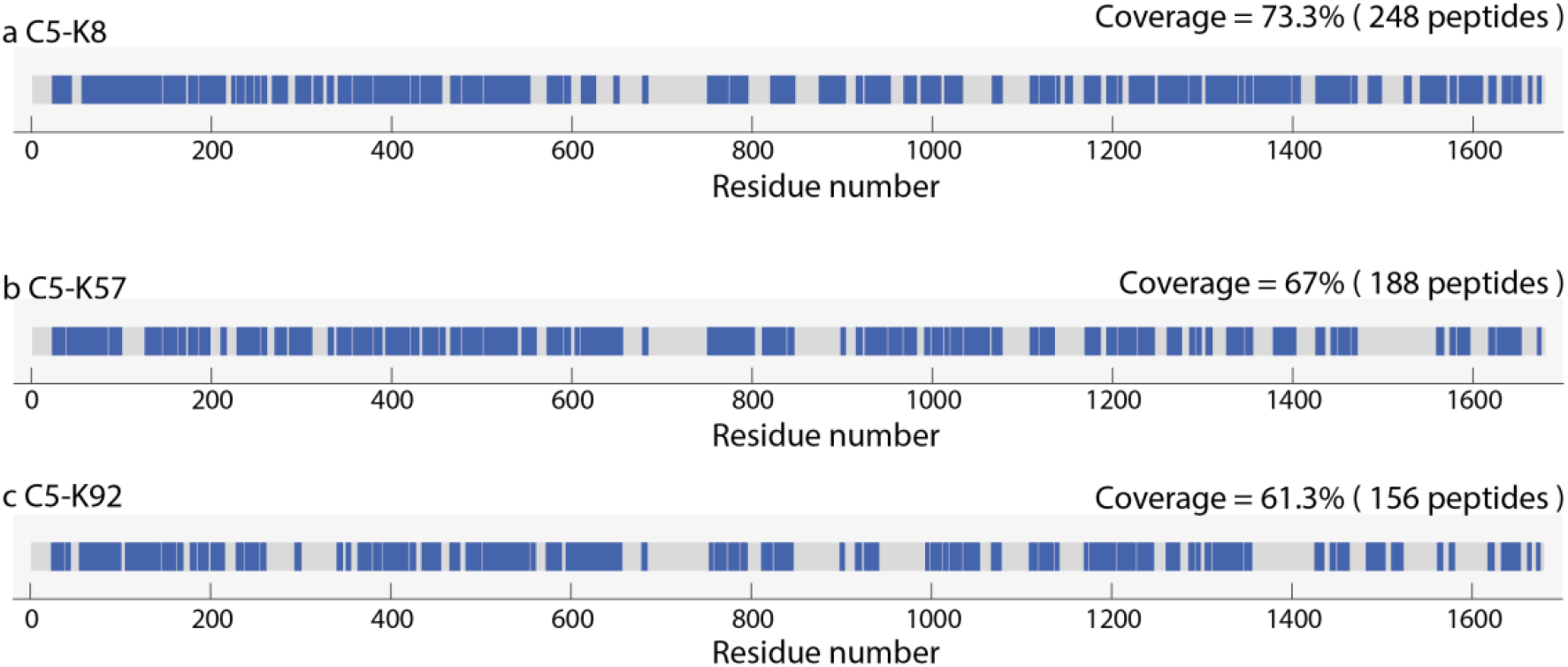
Linear coverage map for C5 data in complex with knob domains

**Supplementary Table 4.3.** HDX data summary table for △HDX of C5 in complex with knob domains K8, K57 and K92.

| Data set | ΔHDX = (C5 + K8) – (C5) | ΔHDX = (C5 + K57) – (C5) | ΔHDX = (C5 + K92) – (C5) |
| --- | --- | --- | --- |
| Protein buffer conditions | PBS, pH 7.2 |  |  |
| Deuterium time course analysed | 30, 120, 900 and 3600 sec |  |  |
| Number of peptides | 248 | 188 | 156 |
| Sequence coverage | 73.3 % | 67.0 % | 61.3 % |
| Redundancy | 2.43 | 2.10 | 2.43 |
| Replicates | Triplicates |  |  |
| Average standard deviation | 0.06 | 0.08 | 0.09 |
| Significant differences at 1 hour ΔHDX* | CI 98% = $0 \pm 24$ Da | CI 98% = $0 \pm 33$ Da | CI 98% = $0 \pm 39$ Da |

\* To compare significant differences, a T-test with $\alpha=0.02$ was used. Only peptides which satisfied a ΔHDX confidence interval of 98 % were considered significant.

**Supplementary 5.1.** SPR single-cycle kinetics of knob domains binding to human C5b Data Table (summary of *n=3*)

| | Chi<sup>2</sup><br>(RU<sup>2</sup>) | mean $k_{on}$<br>(1/Ms) | mean $k_{off}$<br>(1/s) | mean $K_D$<br>(M) | Mean<br>stoichiometric<br>ratio | comment |
| --- | --- | --- | --- | --- | --- | --- |
| K8 | - | - | - | - | 0.1 | Non-binding |
| K92 | 0.04 | 1.14E+05 | 6.20E-05 | 5.48E-10 | 0.4 |  |
| K57 | 0.07 | 2.51E+05 | 3.54E-04 | 1.41E-09 | 0.4 |  |
| K149 | 0.03 | 7.49E+05 | 3.76E-03 | 5.02E-09 | 0.4 |  |

**$n=1$**
| | Chi<sup>2</sup><br>(RU<sup>2</sup>) | $k_{on}$<br>(1/Ms) | $k_{off}$ (1/s) | $K_D$ (M) | Rmax<br>(RU) | Stoichiometric<br>ratio | comment |
| --- | --- | --- | --- | --- | --- | --- | --- |
| K8 | - | - | - | - | 2.1 | 0.1 | Non-binding |
| K92 | 0.06 | 1.11E+05 | 5.20E-05 | 4.67E-10 | 5.4 | 0.4 |  |
| K57 | 0.05 | 2.73E+05 | 3.75E-04 | 1.37E-09 | 6.2 | 0.4 |  |
| K149 | 0.03 | 7.56E+05 | 3.65E-03 | 4.83E-09 | 2.8 | 0.3 |  |

**$n=2$**
| | Chi<sup>2</sup><br>(RU<sup>2</sup>) | $k_{on}$ (1/Ms) | $k_{off}$ (1/s) | $K_D$ (M) | Rmax<br>(RU) | Stoichiometric<br>ratio | comment |
| --- | --- | --- | --- | --- | --- | --- | --- |
| K8 | - | - | - | - | 2.4 | 0.1 | Non-binding |
| K92 | 0.03 | 1.10E+05 | 7.68E-05 | 6.99E-10 | 5.2 | 0.4 |  |
| K57 | 0.08 | 2.40E+05 | 3.46E-04 | 1.44E-09 | 8.2 | 0.4 |  |
| k149 | 0.03 | 7.80E+05 | 3.99E-03 | 5.11E-09 | 5.1 | 0.4 |  |

**$n=3$**
| | Chi<sup>2</sup><br>(RU<sup>2</sup>) | $k_{on}$ (1/Ms) | $k_{off}$ (1/s) | $K_D$ (M) | Rmax<br>(RU) | Stoichiometric<br>ratio | comment |
| --- | --- | --- | --- | --- | --- | --- | --- |
| K8 | - | - | - | - | 2.6 | 0.2 | Non-binding |
| K92 | 0.03 | 1.20E+05 | 5.73E-05 | 4.78E-10 | 4.7 | 0.3 |  |
| K57 | 0.08 | 2.38E+05 | 3.42E-04 | 1.43E-09 | 7.9 | 0.4 |  |
| K149 | 0.03 | 7.12E+05 | 3.65E-03 | 5.13E-09 | 5.2 | 0.4 |  |

**Supplementary Figure 5.2.**
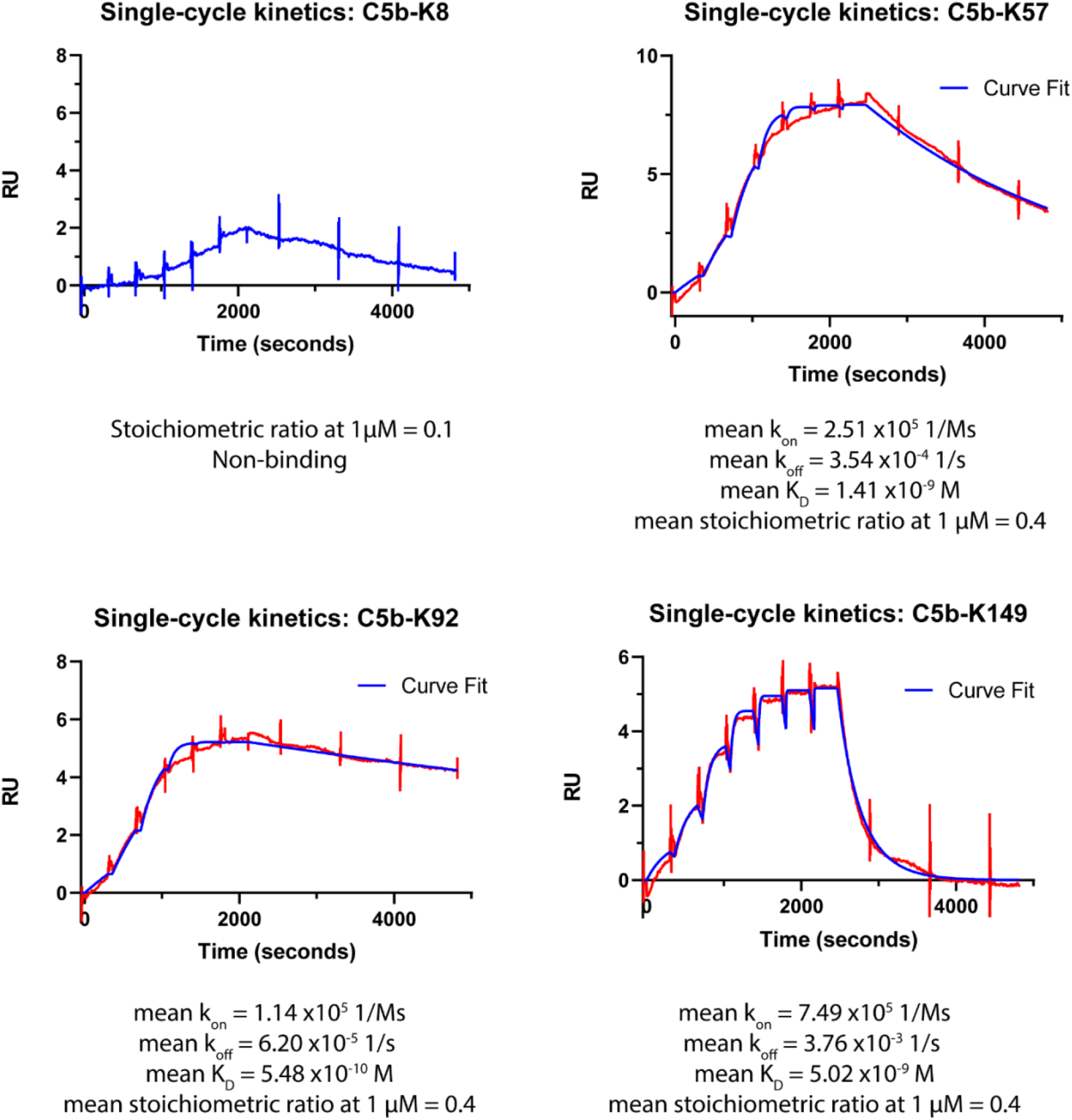
SPR single-cycle kinetics of knob domains binding to human C5b. Example sensorgrams and curve fits (1:1 binding model) are shown with summary kinetics from *n=3* experiments.

